# Mitogen activated protein kinases function as a cohort during a plant defense response

**DOI:** 10.1101/396192

**Authors:** Brant T. McNeece, Keshav Sharma, Gary W. Lawrence, Kathy S. Lawrence, Vincent P. Klink

## Abstract

Mitogen activated protein kinases (MAPKs) play important signal transduction roles. However, little is known regarding whether MAPKs influence the gene expression of other family members and the relationship that expression has to a biological process. Transcriptomic studies have identified MAPK gene expression occurring within root cells undergoing a defense response to a pathogenic event in the allotetraploid *Glycine max*. Furthermore, functional analyses are presented for its 32 MAPKs revealing 9 of the 32 MAPKs have a defense role, including homologs of *Arabidopsis thaliana* MAPK (MPK) MPK2, MPK3, MPK4, MPK5, MPK6, MPK13, MPK16 and MPK20. Defense signal transduction processes occurring through pathogen activated molecular pattern (PAMP) triggered immunity (PTI) and effector triggered immunity (ETI) have been determined in relation to these MAPKs. PTI has been analyzed by examining *BOTRYTIS INDUCED KINASE1* (*BIK1*), *ENHANCED DISEASE SUSCEPTIBILITY1* (*EDS1*) and *LESION SIMULATING DISEASE1* (*LSD1*). ETI has been analyzed by examining the role of the bacterial effector protein harpin and the downstream cell membrane receptor *NON-RACE SPECIFIC DISEASE RESISTANCE1* (*NDR1*). Experiments have identified 5 different types of gene expression relating to MAPK expression. The MAPKs are shown to influence PTI and ETI gene expression and a panel of proven defense genes including an ABC-G type transporter, 20S membrane fusion particle components, glycoside biosynthesis, carbon metabolism, hemicellulose modification, transcription and *PATHOGENESIS RELATED 1* (*PR1*). The experiments show MAPKs broadly influence the expression of other defense MAPKs, including the co-regulation of parologous MAPKs and reveal its relationship to proven defense genes.

## INTRODUCTION

Organisms respond to a number of biotic and abiotic challenges through signal transduction processes, allowing them to cope within their environment. A central signal transduction platform shared between eukaryotes is the three tiered mitogen activated protein kinase (MAPK) cascade (Sturgill and Ray, 1986; Rossomando et al. 1987, 1989; Mohanta et al. 2015). This cascade transduces input information through a stepwise series of phosphorylation events from MAPKKKs (MEKKs), to MAPKKs (MEKs) to MAPKs (MPKs) that leads to an appropriate output response having high fidelity (Jonak et al. 2002; MAPK Group, 2002). In this way, the MAPK platform has been shown to function as a cooperative enzyme functioning in ways that switch cells from one discrete state to another (Huang and Ferrell, 1996; Hazzalin and Mahadevan, 2002). However, in some unusual cases, MPKs can function independently of MEKKs and MEKs through autophosphorylation (Nagy et al. 2015). These studies indicate that much remains to be learned regarding the function of MAPK signaling, particularly in plants.

Early studies of MAPK signaling in plants have benefitted from the sequenced genome of the diploid genetic model *Arabidopsis thaliana* (Tabata et al. 2000; Jonak et al. 2002; MAPK Group, 2002). The genome of *A. thaliana* has 80 MAPKKKs that further transduce signal information through its 10 MAPKKs and then 20 MAPKs, leading to various output responses (Jonak et al. 2002; MAPK Group, 2002). Among the three tiers, much attention has been directed toward the MAPKs and in relation to defense signaling the studies are usually devoted to examining individual or limited numbers of family members (Desikan et al. 1999, 2001; Petersen et al. 2000; Nühse et al. 2000; Asai et al. 2002; Takahashi et al. 2011; Bethke et al. 2012; Eschen-Lippold et al. 2012; Nitta et al. 2014). Other studies have approached studying MAPK signaling in a linear manner from MAPKKK to MAPKK to MAPK (Asai et al. 2002). Studies of the influence between MAPKs in a lateral manner with regard to a whole gene family are absent from the literature.

Regarding plant defense, MAPK signaling has been shown to function downstream of different types of receptor systems that lead to pathogen associated molecular pattern (PAMP) triggered immunity (PTI) and effector-triggered immunity (ETI) (Zhang and Klessig, 2001; Li et al. 2002; Jones and Dangl, 2006; Chinchilla et al. 2007; Lu et al. 2010a, b, Zhang et al. 2010, 2011; Wu et al. 2011; Schwessinger et al. 2011; Chang and Nick 2012; Couto et al. 2016). PTI can be initiated through the toll-interleukin receptor nucleotide binding leucine rich repeat resistance (TIR)-NB-LRR R protein RECOGNITION OF PERONOSPORA PARASITICA 4 (RPP4) (among others) in processes that lead to ENHANCED DISEASE SUSCEPTIBILITY1 (EDS1)-driven engagement of defense gene expression that may or may not also be accompanied by salicylic acid (SA) signaling (Aarts et al. 1998; Wildermuth et al. 2001; van der Biezen et al. 2002; Wiermer et al. 2005; Kesarwani et al. 2007; Tsuda et al. 2008; Vlot et al., 2009; Huang et al. 2010; Eitas and Dangl 2010; Gouhier-Darimont et al. 2013; Hilfiker et al. 2014; Zhang et al. 2017). ETI can be initiated by the bacterial effector harpin that engages the transcription of the coiled-coil nucleotide binding leucine rich repeat (CC-NB-LRR) NON-RACE SPECIFIC DISEASE RESISTANCE 1 (NDR1)/HARPIN INDUCED1 (HIN1) (Wei et al. 1992; Century et al. 1995, 1997; Gopalan et al. 1996). NDR1 then transduces the signal through the MAPK cascade to elicit a defense response (Desikan et al. 1999; 2001; Lee et al. 2001).

Complicating the understanding of these receptor systems are other proteins that have roles that appear to be protective in nature, mitigating the action of pathogen effectors while allowing their partnered R protein to drive defense processes. In some of these cases, the pathogen-inactivated protein elicits a type of autoimmune response that can be highly deleterious to the normal growth of the plant. For example, the PTI protein RPP4/CHILLING-SENSITIVE2 (CHS2) is the paralog of RPP5/SUPPRESSOR OF npr1-1, CONSTITUTIVE 1 (SNC1) (Li et al. 2001; van der Bizen et al. 2002; Huang et al. 2010a). While both RPP4 and RPP5 function through EDS1 to drive defense gene expression, only RPP5 functions along with SA (van der Bizen et al. 2002; Huang et al. 2010a). RPP4 and RPP5 also function along with other TIR-NB-LRR class genes in complex ways as revealed by mutant analyses examining autoimmunity (Chung et al. 2013; Roberts et al. 2013; Dong et al. 2016). For example, the CALMODULIN-BINDING TRANSCRIPTION ACTIVATOR 3 (CAMTA3) appears to be protected by TIR-NB-LRR DOMINANT SUPPRESSOR OF CAMTA3 NUMBER 1 (DSC1) and DSC2, leading to PTI (Bjornson et al. 2014; Rahman et al. 2016; Lolle et al. 2017). Similarly, the ETI protein NDR1 appears to be protected by the unstructured protein RPM1-INTERACTING PROTEIN 4 (RIN4) and two CC-NB-LRR R proteins including RESISTANCE TO PSEUDOMONAS SYRINGAE2 (RPS2) and RESISTANCE TO PSEUDOMONAS SYRINGAE PV MACULICOLA1 (RPM1) (Kunkel et al. 1993; Grant et al. 1995; Mackey et al. 2002, 2003; Axtell and Staskawicz, 2003; Belkhadir et al. 2004; Kim et al. 2005; Day et al. 2006). In related experiments, cross-talk has been shown to occur between PTI and ETI receptor systems (van der Biezen et al. 2002; Veronese et al. 2006; Thomma et al. 2006; Zipfel et al. 2006; Liu et al. 2013; Lolle et al. 2017; Jacob et al. 2018). Consequently, more study is required to understand the fine details of these receptor systems in relation to defense processes to different pathogens and also the MAPK gene family since it functions downstream and acts as an apparent convergence point for ETI and PTI.

An important question that has not received much attention is whether and if so how extensively MAPK gene family members cross-talk or function coordinately on a tier-wide basis during a biological process. For example, work on defense to pathogens in *A. thaliana* has had most of its attention focused on MPK3 and/or its paralog MPK6, with less work done on MPK2 and MPK4 (Petersen et al. 2000; Nühse et al. 2000; Asai et al. 2002). Other studies have examined MPK7, MPK8, MPK9 and MPK12 (Doczi et al. 2007; Takahashi et al. 2011; Jammes et al. 2011). Similar approaches have been made in important agricultural crops such as *Glycine max* (soybean) (Liu et al. 2011, 2014). Experiments performed in transgenic *A. thaliana* have also expressed epitope tagged versions of pathogen elicitors to determine their impact on MAPK activity. For example, transgenic *A. thaliana* expressing the bacterial flg22 elicitor results in phosphorylation of MPK1, MPK11 and MPK13, indicating very specialized roles for these proteins (Bethke et al. 2012; Eschen-Lippold et al. 2012; Nitta et al. 2014). However, a biological function for these activated MAPKs had not been presented. Without a functional characterization, much of these and related expression studies on the MAPK gene family remain uncharacterized observations (Reyna et al. 2006; Lee et al. 2008; Yang et al. 2013; Neupane et al. 2013; Nitta et al. 2014; Asif et al. 2014; Lu et al. 2015).

Employing transgenic approaches similar to those used to identify genes functioning in defense, an analysis is presented here examining each of the 32 members of the MAPK gene family of *G. max* in relation to its defense response during infection by a root pathogen (Bartsch et al. 2006; Klink et al. 2007, 2010, 2011; Schmutz et al. 2010; Grant et al. 2010; Liu et al. 2011; Goodstein et al. 2012; Matsye et al. 2012; Neupane et al. 2013; Mohanta et al. 2015; McNeece et al. 2017). The *G. max* experimental system presents advantages over *A. thaliana* where a number of the defense genes have pleiotropic effects in single or combinatorially-mutated genetic backgrounds that result in highly compromised or lethal phenotypes (Lukowitz et al. 1996; Heese et al. 2001; Collins et al. 2003; Assaad et al. 2004; Zhang et al. 2007; Lipka et al. 2008; Han et al. 2010; Humphry et al. 2010; Huang et al. 2010a; Heidrich et al. 2011; Johansson et al. 2014). Experiments presented here show that *G. max* MAPKs (MAPKs) identified to be expressed within parasitized root cells undergoing the process of defense to the root pathogenic nematode *Heterodera glycines* often presage their function in defense. Experiments show the relationship between PTI and ETI and the defense MAPKs to prior published work (Matsye et al. 2011, 2012; Pant et al. 2014, 2015; Sharma et al. 2016; McNeece et al. 2017; Aljaafri et al. 2017; Klink et al. 2017). As presented in earlier experiments, PTI has been studied here by examining *ENHANCED DISEASE SUSCEPTIBILITY1* (*EDS1*) and *LESION SIMULATING DISEASE1* (*LSD1*) (Dietrich et al. 1997; Aarts et al. 1998; Falk et al. 1999; Pant et al. 2014, 2015; Matuszkiewicz et al. 2018). ETI has been studied by examining the relationship between treatment with bacterial elicitor harpin and MAPK gene expression (Aljaafri et al. 2017). The experiments presented here then focus on *G. max* homologs of the ETI genes *NON-RACE SPECIFIC DISEASE RESISTANCE 1/HARPIN INDUCED1* (*NDR1/HIN1*) and *BOTRYTIS INDUCED KINASE1* (*BIK1*) (Wei et al. 1992; Century et al. 1995, 1997; Gopalan et al. 1996; Veronese et al. 2006; McNeece et al. 2017; Aljaafri et al. 2017). Furthermore, transcriptional analyses show that the transgenic MAPK lines affect the expression of different suites of proven defense genes (Matsye et al. 2012; Pant et al. 2014; Sharma et al. 2016; Klink et al. 2017; McNeece et al. 2017). These genes include those composing the PTI and ETI signal transduction cascades, the 20S membrane fusion particle, alpha hydroxynitrile biogenesis and cyanide metabolism, an ABC-G type transporter, carbon and hemicellulose metabolism and the expression of *PATHOGENESIS RELATED1* (*PR1*) (Matsye et al. 2011, 2012; Pant et al. 2014; Sharma et al. 2016; McNeece et al. 2017; Klink et al. 2017). To our knowledge, this analysis represents the first functional examination of an entire MAPK gene family in any plant species. In the process of the experiments the work is demonstrating that MAPKs exhibit a broad and possibly coordinated influence on the expression of other MAPKs and proven defense genes as it overcomes pathogen infection. Furthermore, the work provides a framework for related experimentation to understand other aspects of MAPK signaling in plants.

## MATERIALS and METHODS

### Identification of *G. max* MAPK genes

*G. max* can become parasitized by the root pathogenic nematode *H. glycines*, having a life cycle of approximately 30 days (**Figure 1**) (Lauritis et al. 1983). In prior experiments two different *H. glycines*-resistant genotypes *G. max*_[Peking/PI 548402]_ and *G. max*_[PI 88788]_ have been infected with *H. glycines*_[NL1-Rhg/HG-type 7/race 3]_, resulting in the expected resistant outcome (Ross 1958; Endo 1965, 1991; Klink et al. 2007, 2009a, 2010a, b, 2011; Matsye et al. 2011). Laser capture microdissection (LCM) has been used to collect pericycle cells at a time point prior to *H. glycines* infection (the 0 days post infection [dpi] control) and syncytia undergoing the process of resistance at 3 and 6 dpi (Klink et al. 2007, 2009a, 2010a, b, 2011; Matsye et al. 2011). Affymetrix^®^ Soybean Gene Chip^®^ hybridizations to produced cDNA probe synthesized from RNA isolated from these samples have been run in triplicate, resulting in the production of 6 total arrays for each time point (*G. max*_[Peking/PI 548402]_: arrays 1-3; *G. max*_[PI 88788]_: arrays 1-3) (**Supplemental Table 1**). The gene expression analysis procedure has used the Bioconductor implementation of the standard Affymetrix^®^ detection call methodology (DCM). MAPK gene expression data that has been generated by LCM-DCM has been extracted from those analyses and used in the presented study. Additional gene expression data for a family of *G. max* pathogenesis related 1 (GmPR1) genes, including GmPR1-6 (Glyma15g06780) has been extracted. A cutoff has been set whereby a gene is considered expressed in a given sample if probe signal is measurable (M) above threshold (p < 0.05) on all 6 arrays (combining the 3 arrays from *G. max*_[Peking/PI 548402]_ and *G. max*_[PI 88788]_ according to the described methods (**Supplemental Table 1**) (Klink et al. 2007, 2009a, 2010a, b, 2011; Matsye et al. 2011). The expression of a gene has been considered not measured (N/M) if probe signal is not detected at a statistically significant level (p ≥ 0.05) on any one of the 6 total arrays. Due to how the Affymetrix ^®^ microarray has been constructed, some genes have had no probe set fabricated onto the array. In these cases, gene expression could not be quantified according to the analysis procedures so the gene expression analysis is not applicable (n/a) to those genes. Consequently, that characteristic of the Affymetrix^®^ Soybean Gene Chip^®^ influenced the decision to produce the comprehensive analysis of the *G. max* MAPK gene family. However, gene expression data generated by RNA seq examining RNA isolated from whole root libraries or microarrays has been obtained for those MAPKs genes functioning in defense, but whose gene expression could not be determined (n/a) in our earlier studies (Goodstein et al. 2012; Neupane et al. 2013).

**Figure 1.**
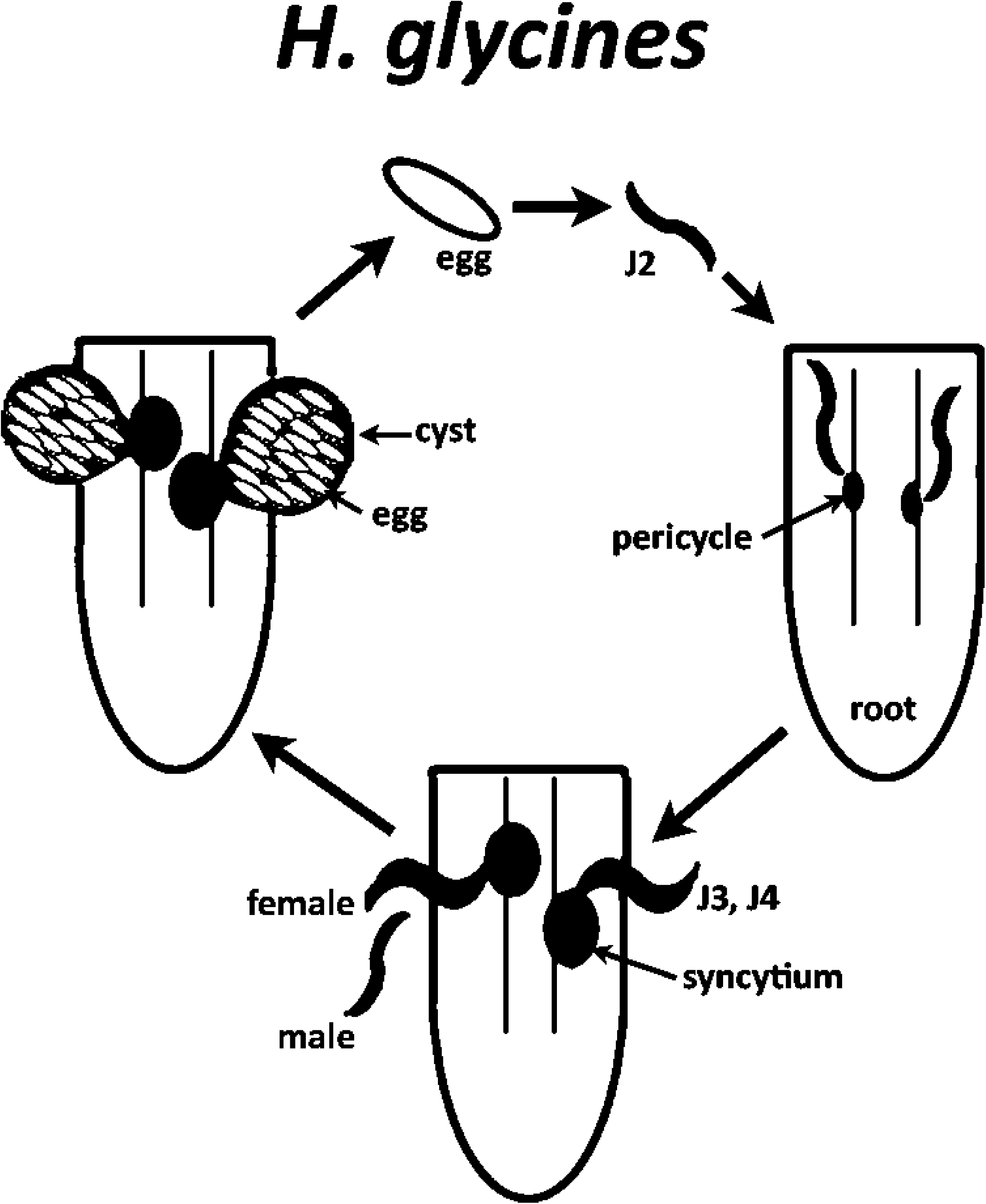
*H. glycines* life cycle. Egg; juvenile stages, second (J2), third (J3), fourth (J4). Male, female; parasitized pericycle cells develop into a syncytium which is a nursing structure composed from the merged cytoplasm of 200-250 cells that also serves as the site of the localized defense response; cyst, female carcass structure containing the eggs.

### Cloning and production of genetic constructs

The *G. max* genome has undergone three different analyses examining the breadth of the MAPK gene family (Liu et al. 2011; Neupane et al. 2013; Mohanta et al. 2015). Liu et al. (2011) stated there are at least 56 MAPKs that had been identified by reciprocal BLASTN searches between NCBI and the *G. max* genome housed at Phytozome (Goodstein et al. 2012). Neupane et al. (2013), employing the hidden Markov model (HMM) for their analysis, stated there are 38 *G. max* MAPKs, including a new clade E containing 6 MAPKs not present in other flowering plants. In contrast, Mohanta et al. (2015) used stricter criteria, resulting in the identification of 31 MAPKs belonging to the canonical clades A-D found in *A. thaliana*. Analyses show that the clade E MAPKs of Neupane et al. (2013) are not closely related to *A. thaliana* MPKs (**Supplemental Table 2**) (Mohanta et al. 2015). Consequently, the analysis presented here has been limited to the MAPK gene family identified by Mohanta et al. (2015) with the addition of MAPK16-4 (Glyma07g38510) identified in Neupane et al. (2013) which clearly is a MAPK, but has somehow remained absent in the analysis of Mohanta et al. (2015). The MAPK numbering scheme of Neupane et al. (2013) has been used in the analysis presented here because it relates more closely to the corresponding *A. thaliana* MPKs (MAPK Group, 2002). The MAPK gene sequences have been downloaded from the *G. max* genome database housed at Phytozome (Goodstein et al. 2012). Cloning of the MAPK genes have been accomplished to permit engineering their overexpression (OE) in the *H. glycines*-susceptible *G. max*_[Williams 82/PI 518671]_ or RNA interference (RNAi) in the *H. glycines*-resistant *G. max*_[Peking/PI 548402]_ (Klink et al. 2009a; Matsye et al. 2012). The PCR and quantitative PCR (qPCR) primers are provided (**Supplemental Table 3**).

### Cloning MAPKs into the pRAP destination vectors

The pRAP destination vectors are based off of the published Gateway^®^ cloning vector platform (Curtis and Grossniklaus 2003; Klink et al. 2009b; Matsye et al. 2012). The MAPKs, have been ligated in the pENTR/D-TOPO^®^ entry vector (Invitrogen^®^) according to the manufacturer’s instructions. LR Clonase^®^ (Invitrogen^®^) has been used to facilitate the transfer the MAPK amplicon to the pRAP15 overexpression and pRAP17 RNAi destination vectors according to the manufacturer’s instructions. The unengineered pRAP15 or pRAP17 control has the *ccd*B gene located in the position where, otherwise, the MAPK amplicon is inserted during the LR clonase reaction. This feature makes the pRAP15-*ccd*B (overexpression control) and pRAP17-*ccd*B (RNAi control) unengineered vectors suitable controls for non-specific effects caused by gene overexpression or RNAi (Klink et al. 2009b; Matsye et al. 2012; Pant et al. 2014; Sharma et al. 2016; McNeece et al. 2017). The pRAP15-*ccd*B, pRAP15-MAPK-OE, pRAP17-*ccd*B and pRAP17-MAPK-RNAi vectors have been used in freeze-thaw incubation experiments to transform chemically competent *Agrobacterium rhizogenes* K599 (K599) with the designated gene cassette (Hofgen and Willmitzer 1988; Haas et al. 1995; Collier et al. 2005; Klink et al. 2009b).

### Production of genetically mosaic transgenic *G. max* plants

The enhanced green fluorescent reporter (eGFP) has been used for visual selection of transgenic *G. max* roots (Haseloff et al. 1997; Klink et al. 2009b; Matsye et al. 2012). Roots exhibiting the eGFP reporter expression will also possess the MAPK expression cassette, each with their own promoter and terminator sequences (Klink et al. 2009b; Matsye et al. 2012). K599 transfers the DNA cassettes located between the left and right borders of the destination vector into the root cell chromosomal DNA, resulting in stable transformation event in somatic cells even though the construct is not incorporated into the germline. The transformed cell subsequently develops through mitotic and developmental events into a transgenic root from the base of the shoot stock (**Supplemental Figure 1**) (Klink et al. 2008). The resultant plant is a genetic mosaic called a composite plant with a non-transgenic shoot having a transgenic root system (Tepfer, 1984; Haas et al. 1995; Collier et al. 2005; Klink et al. 2009b; Matsye et al. 2012). Consequently, each individual transgenic root system is an independent transformant line (Tepfer, 1984; Klink et al. 2009b; Matsye et al. 2012; Matthews et al. 2013). The transgenic plants are planted in SC10 Super cone-tainers secured in RL98 trays and recover for two weeks prior to experiments (Stuewe and Sons, Inc.^®^) (McNeece et al. 2017). The functionality of the genetic constructs in *G. max* is confirmed by qPCR (Please refer to quantitative PCR [qPCR] section).

### cDNA synthesis

*G. max* root mRNA has been isolated according to our published procedures using the UltraClean^®^ Plant RNA Isolation Kit (Mo Bio Laboratories^®^, Inc.) according to the manufacturer’s instructions (Matsye et al. 2012). Genomic DNA has been removed from the mRNA with DNase I (Invitrogen^®^) according to the manufacturer’s instructions. The cDNA has been synthesized from mRNA using the SuperScript^®^ First Strand Synthesis System for RT-PCR (Invitrogen^®^) using the oligo d(T) as the primer according to the manufacturer’s instructions. Genomic DNA contamination has been assessed by our published methods by using β-conglycinin primer pair that amplifies DNA across an intron, yielding different sized products based on the presence or absence of that intron (Klink et al. 2009b).

### Assessment of gene expression by qPCR

Assessment of engineered MAPK expression in *G. max* and an examination of proven defense gene expression has been accomplished by qPCR using Taqman^®^ 6-carboxyfluorescein (6-FAM) probes and Black Hole Quencher (BHQ1) (MWG Operon; Birmingham, AL) (**Supplemental Table 3**) (Sharma et al. 2016). The qPCR control that has been used in the *G. max* experiments has been designed from an expressed sequence tag of ribosomal S21 protein coding gene which functions like other proven control genes (Klink et al. 2005; Sharma et al. 2016). The fold change in gene expression caused by the genetic engineering event has been calculated using 2^-ΔΔ*C*^_T_ according to our prior published methods (Livak and Schmittgen 2001; McNeece et al. 2017).

### Analysis of the effect MAPK genetic constructs have on *H glycines* parasitism in *G. max*

*glycines* cysts have been collected from transgenic roots according to our procedures (**Supplemental Figure 1**) (McNeece et al. 2017). An analysis of the effect that the MAPK genetic constructs have on *H glycines* parasitism in *G. max* has been calculated and presented as the female index (FI), which is the community-accepted standard representation of the obtained data (Golden et al. 1970). The FI = (Nx/Ns) X 100, where Nx is the average number of females on the test cultivar and Ns is the average number of females on the standard susceptible cultivar (Golden et al. 1970). Using the formula of Golden et al. (1970), Nx is the pRAP15-MAPK or pRAP17-MAPK-transformed line and Ns is the pRAP15-*ccd*B or pRAP17-*ccd*B control line which has been calculated according to our prior published methods (Matsye et al. 2012; Pant et al. 2014, 2015; Sharma et al. 2016; McNeece et al. 2017). The FI has been calculated as cysts per mass of the whole root (wr) and also cysts per gram (pg) of root according to our published methods (Pant et al. 2015; Sharma et al. 2016; McNeece et al. 2017; Klink et al. 2017). The whole root analysis is how the data has been presented historically (Golden et al. 1970). The cyst per gram analysis accounts for possible altered root growth caused by the influence of the overexpression or RNAi of MAPK (or GmPR1-6). Three biological replicates have been made for each construct with 10-20 individual transgenic plants each. A statistical analysis has been performed using the Mann–Whitney–Wilcoxon (MWW) Rank-Sum Test, p < 0.05 cutoff (Mann and Whitney, 1947; McNeece et al. 2017). The MWW Rank-Sum Test, as stated, is a nonparametric test of the null hypothesis not requiring the assumption of normal distributions (Mann and Whitney, 1947).

### Microscopy

Stereoscope images have been obtained on a Wild Heerbrugg stereoscope. The lenses are Wild Heerbrugg Makrozoom 1:5 having a 6.3-32x scale. Image capture has been done using the IMT i-solution computer package (IMT i-solution Inc., Ho Chi Minh City, Vietnam).

### Bioinformatics

Signal peptide prediction has been determined using SignalP-4.1, Phobius and PrediSI prediction (Hiller et al. 2004; Käll et al. 2007; Petersen et al. 2011). The analysis has been performed on the GmPR1-6 protein sequence. The analysis has been performed to determine how GmPR1-6 would relate to the vesicle-mediated secretion system, a cellular apparatus whose components have been shown to function in defense in this pathosystem (Matsye et al. 2012; Pant et al. 2014; Sharma et al. 2016).

## RESULTS

### Identification members of the *G. max* MAPK gene family

The analysis presented here aims at understanding the expression of the *G. max* MAPK (GmMAPK) gene family as it relates to their function during a pathogenic process (**Figure 2**). *G. max* has 32 MAPKs exhibiting clear phylogenetic relationships with those found in *A. thaliana* (**Supplemental Table 1**) (Neupane et al. 2013; Mohanta et al. 2015). Six additional kinases rooted at the base of the MAPK phylogenetic tree are not similar to the *A. thaliana* MPKs (**Supplemental Table 2**) (Arabidopsis Interactome Mapping Consortium, 2011; Neupane et al. 2013; Nemoto et al. 2015; Mohanta et al. 2015). Consequently, the analysis of those genes are beyond the scope of this presented study, justifying the selection of the 32 MAPKs studied herein.

**Figure 2.**
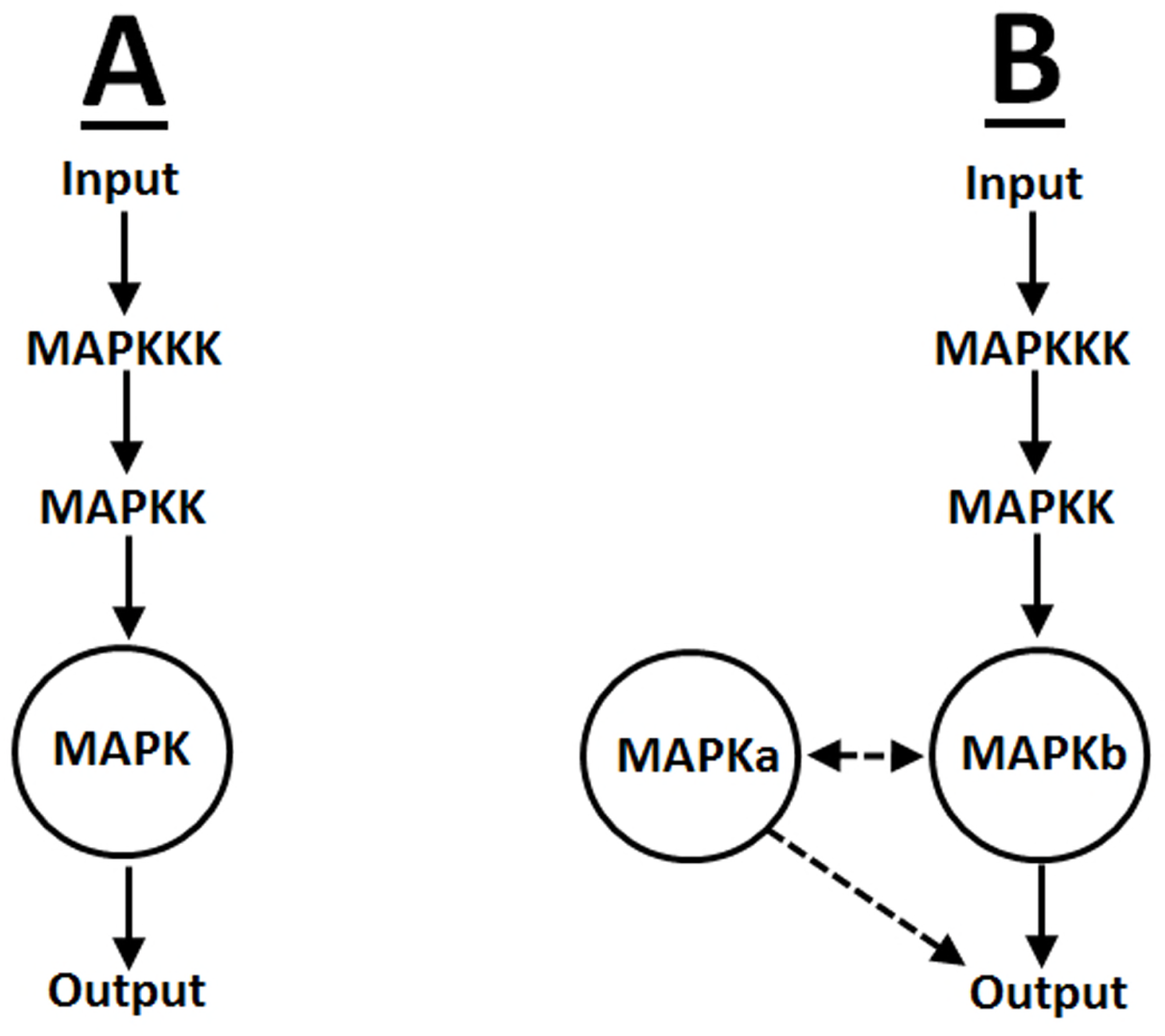
MAPK models. **A**. An input signal is transduced through the three-tiered MAPK pathway, engaging an output response (e.g. defense). **B**. An input signal is transduced through the MAPK pathway with crosstalk (dashed arrows) occurring at the MAPK tier between different MAPKs (e.g. MAPKa, MAPKb), eliciting an output response.

### *G. max* MAPKs are expressed within root cells undergoing a defense response

A transcript analysis examining MAPK expression occurring within syncytia undergoing the process of defense in two different *H. glycines*-resistant genotypes has been performed (**Table 1; Supplemental Table 1**). The experiments have identified the expression of 12 MAPKs occurring in pericycle and/or syncytia undergoing processes normally leading to a defense response. The remaining MAPKs either did not exhibit measurable expression or their transcript abundance could not be assessed due to how the Affymetrix^®^ Gene Chip^®^ had been fabricated. Many of the MAPKs whose expression could not be assessed here have been shown to be expressed within roots (Neupane et al. 2013). These MAPKs include MAPK1, MAPK3-2, MAPK4-2, MAPK6-1, MAPK9-1, MAPK11-2, MAPK20-1 and MAPK20-3. In three cases, MAPK5-3, MAPK9-3 and MAPK9-4, expression has not been identified in root tissue in Neupane et al. (2013). However, RNA seq data has identified MAPK5-3, MAPK9-3 and MAPK9-4 to be expressed within roots, lateral roots, root tips and root hairs (Goodstein et al. 2012). The observed expression of each MAPK in roots has influenced the decision to produce the comprehensive analysis of the *G. max* MAPK gene family presented here. Part of this reason is that MAPK expression occurring outside the syncytium could be important to defense processes when considering the possibility that the root may detect the production of damage trails made by the nematode as it is burrowing toward the pericycle cells as early as 12 hours post infection before the production of syncytia can even happen (Klink et al. 2007b).

**Table 1.** LCM-DCM gene expression summary of studied MAPKs (Details are provided in **Supplemental Table 2**). *, the MAPK is proven to have a role in defense. Group (A-D) refers to the different groups of MAPKs that have been classified based on activation loop domain composition (MAPK Group, 2002; Andreasson and Ellis, 2009). M, gene expression is measured as determined by the set criteria of a p < 0.05 in all 6 arrays; N/M, gene expression is not measured as determined by the set criteria since at least one of the 6 arrays had a p ≥ 0.05; n/a, not applicable since the Affymetrix^®^ microarray did not have a probe set fabricated for that gene.

|  |  |  | Time point (dpi) |  |  |
| --- | --- | --- | --- | --- | --- |
| Gene name | Genome accession | Group | 0 | 3 | 6 |
| GmMAPK 1 | Glyma04g03210 | A | n/a | n/a | n/a |
| GmMAPK 2* | Glyma06g03270 | A | N/M | N/M | M |
| GmMAPK 3-1* | Glyma11g15700 | B | M | M | M |
| GmMAPK 3-2* | Glyma12g07770 | B | n/a | n/a | n/a |
| GmMAPK 4-1* | Glyma07g07270 | C | M | M | M |
| GmMAPK 4-2 | Glyma16g03670 | C | n/a | n/a | n/a |
| GmMAPK 5-1 | Glyma01g43100 | C | N/M | N/M | M |
| GmMAPK 5-2 | Glyma11g02420 | C | N/M | N/M | N/M |
| GmMAPK 5-3* | Glyma08g02060 | C | n/a | n/a | n/a |
| GmMAPK 5-4 | Glyma05g37480 | C | N/M | N/M | N/M |
| GmMAPK 6-1 | Glyma07g32750 | B | n/a | n/a | n/a |
| GmMAPK 6-2* | Glyma02g15690 | B | M | M | M |
| GmMAPK 7 | Glyma05g28980 | A | N/M | N/M | N/M |
| GmMAPK 9-1 | Glyma05g33980 | D | n/a | n/a | n/a |
| GmMAPK 9-2 | Glyma08g05700 | D | N/M | N/M | N/M |
| GmMAPK 9-3 | Glyma07g11470 | D | n/a | n/a | n/a |
| GmMAPK 9-4 | Glyma09g30790 | D | n/a | n/a | n/a |
| GmMAPK 11-1 | Glyma18g47140 | C | N/M | M | M |
| GmMAPK 11-2 | Glyma09g39190 | C | n/a | n/a | n/a |
| GmMAPK 13-1* | Glyma12g07850 | B | N/M | M | M |
| GmMAPK 13-2 | Glyma11g15590 | B | N/M | N/M | N/M |
| GmMAPK 14 | Glyma08g12150 | A | N/M | N/M | N/M |
| GmMAPK 16-1 | Glyma13g28120 | D | N/M | N/M | N/M |
| GmMAPK 16-2 | Glyma15g10940 | D | N/M | M | M |
| GmMAPK 16-3 | Glyma17g02220 | D | N/M | M | M |
| GmMAPK 16-4* | Glyma07g38510 | D | M | M | M |
| GmMAPK 18 | Glyma15g38490 | D | N/M | N/M | N/M |
| GmMAPK 19 | Glyma13g33860 | D | N/M | N/M | M |
| GmMAPK 20-1 | Glyma18g12720 | A | n/a | n/a | n/a |
| GmMAPK 20-2* | Glyma14g03190 | A | N/M | M | N/M |
| GmMAPK 20-3 | Glyma02g45630 | A | n/a | n/a | n/a |
| GmMAPK 20-4 | Glyma08g42240 | A | N/M | N/M | N/M |

### Characterization of the *G. max* MAPK gene family as it relates to defense

Functional, transgenic experiments have been designed to examine if any of the 32 *G. max* MAPKs act during the root defense process to *H. glycines* parasitism. Each of the 32 MAPKs have been examined in overexpression and RNAi experiments. By performing the experiments in this manner, a comprehensive understanding of the involvement of the MAPKs in defense will have been ascertained.

Overexpression experiments have increased the relative transcript abundance of each of the 32 individual MAPKs (**Supplemental Figure 2**). Prior to performing the experiments it had been noted that transgenically-driven target gene overexpression could (1) have no effect, (2) increase or (3) decrease root mass (**Figure 3**). Consequently, the analysis methods have measured the female index (FI) in relation to the whole root mass and then standardized those data by also calculating cysts per gram of root mass. The results of the analyses have identified a statistically significant decrease in parasitism in both the wr and pg analyses in 15 of the replicated transgenic MAPK-overexpression experiments (**Figure 3**). Notably, a number of wr analyses have identified MAPK-OE lines meeting the criteria for suppressed *H. glycines* parasitism. However, these results contrast with the outcome having been obtained in pg analyses showing a statistically significant difference (increase) in FI. In these cases, the results indicate that the transgenic event has a negative effect on root growth (**Supplemental Figure 3**). Of the 15 OE MAPKs that decrease nematode parasitism in both the wr and pg analyses, MAPK2 and MAPK5-3 influence root growth (**Supplemental Figure 3**). In each case, root mass is decreased. For the remaining 13 OE MAPKs, their overexpression has not been observed to impact root mass to a statistically significant level under the analysis procedures (**Supplemental Figure 3**). However, other MAPKs have been shown to decrease root mass (**Supplemental Figure 3**). These MAPKs include MAPK4-2, MAPK5-1, MAPK7, MAPK11-2, MAPK13-2, MAPK16-1 and MAPK19. Under the set criteria, in no case have observations been made obtaining a statistically significant increase in root mass. However, an increase in root mass has been observed in two of the three replicates for the defense MAPK6-2 and the non-defense MAPK9-3 and MAPK9-4 (**data not presented**).

**Figure 3.**
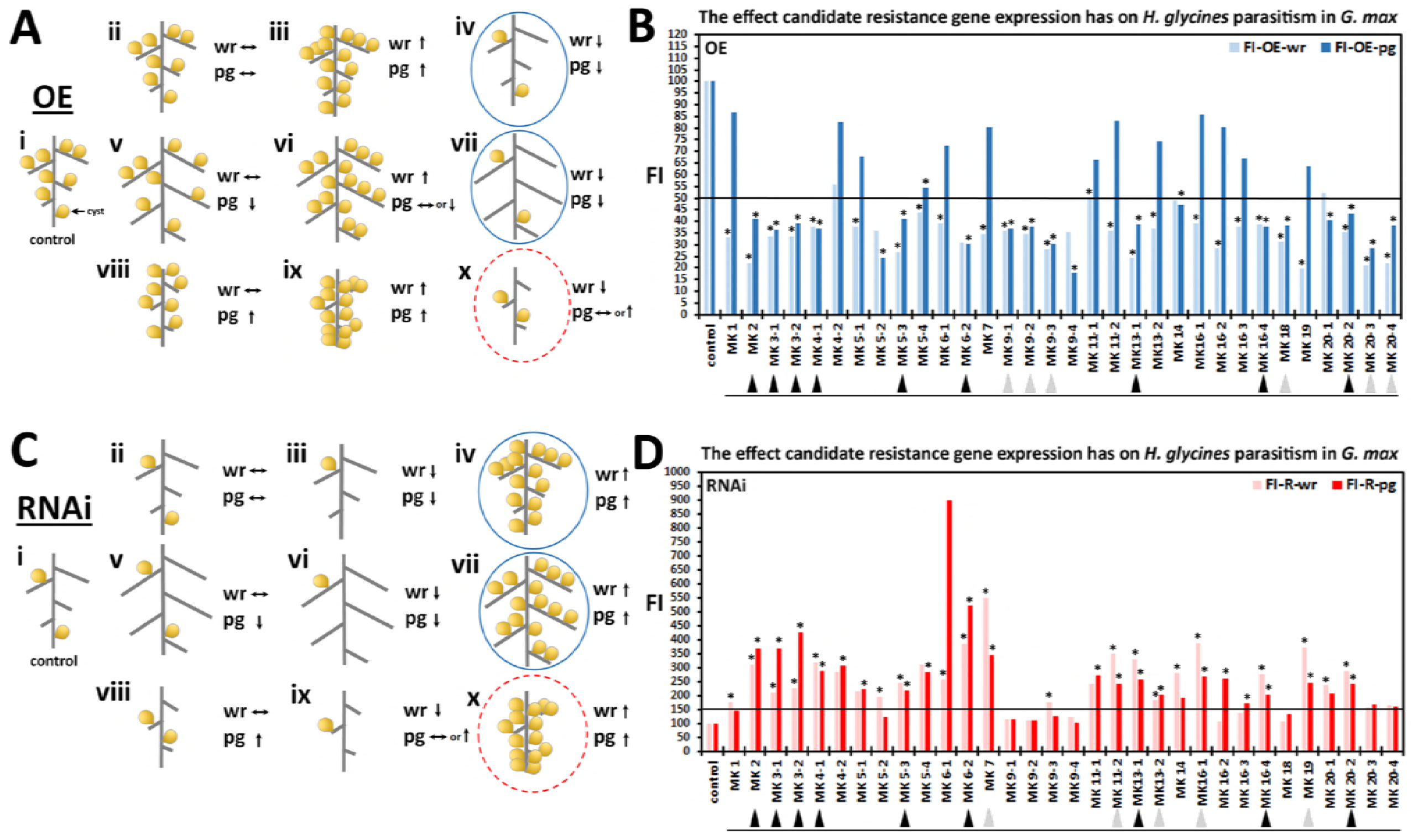
Experimental analysis of MAPK function in defense as presented by the FI. **A**. Possible OE outcomes, i, control plants have a defined root mass and level of nematode parasitism; ii-iv, the overexpressed MAPK has no effect on root mass as compared to the control; v-vii, the overexpressed MAPK increases root mass as compared to the control; viii-x, the overexpressed MAPK decreases root mass as compared to the control. Parasitism is measured in two different manners, cysts per whole root (wr) and also standardized per gram (pg). ↔, no change in parasitism as compared to the control; ↑, an increase in parasitism as compared to the control; ↓ a decrease in parasitism as compared to the control. The blue circles indicate the desired outcomes whereby the expression of the genetic element either has no deleterious effect on root mass or increases root mass as compared to the control. The red dashed circles indicate the undesired outcomes whereby the expression of the genetic element has a deleterious effect on root mass as compared to the control. **B**. FI in OE lines. The lighter blue histograms are measurements per wr. The darker blue histograms are measurements pg. The black bar indicates the cutoff of a FI < 50. * p < 0.05 calculated by the Mann–Whitney–Wilcoxon (MWW) Rank-Sum Test which is a nonparametric test of the null hypothesis not requiring the assumption of normal distributions (Mann and Whitney, 1947). The experimental error representing standard deviation is presented. The gray and black triangles present below the MAPK identifiers on the x-axis represent genes whose overexpression has a significant decrease in FI (y-axis) and were statistically significant in the wr and pg analyses. The black triangles (n = 9) denote genes that also exhibited a statistically significant reduction in the RNAi lines (see below). **C**, Possible RNAi outcomes with description the same as presented in **A**. **D**. FI in RNAi lines. The lighter red histograms are measurements per wr. The darker red histogram are measurements pg. * p < 0.05. The black bar indicates the cut off of a FI > 150. The experimental error representing standard deviation is presented. Please refer to Methods for analysis details.

RNAi experiments have decreased the relative transcript abundance of each of the 32 individual MAPKs (**Supplemental Figure 2**). The RNAi of an examined gene can also have no effect, increase or decrease root mass (**Figure 3**). Consequently, the effect that MAPK-RNAi transgene expression exerts on root development has been considered in the analysis procedures. The results of the analyses have identified a statistically significant increase in parasitism in both the wr and pg analyses in 14 of these replicated transgenic experiments (**Figure 3**). A number of wr analyses have identified MAPK-RNAi lines meeting the criteria for suppressed *H. glycines* parasitism. However, these results have contrasted with the outcome obtained in pg analyses showing a statistically significant difference (decrease) in FI. In these cases the results have identified a negative effect on root mass. However, in three cases (e.g. MAPK3-1, MAPK3-2 and MAPK6-2) the FI increased even further in the RNAi lines at a statistically significant level, possibly indicative of impaired root growth. In fact, a number of MAPKs are observed to exhibit variations in FI between the wr and pg analyses, indicating the transgenically-expressed RNAi construct influences root growth (**Supplemental Figure 3**). The RNAi of MAPK3-2 has been observed to decrease root mass while the RNAi of MAPK4-1 increases root mass. For the remaining defense MAPKs, their RNAi has not been observed to impact root mass to a statistically significant level. No statistically significant change in root mass has been observed for the remaining non-defense MAPK RNAi lines.

### Congruence exists between the MAPK overexpression and RNAi analyses

Analyses have then been performed to determine if the same MAPKs that affect *H. glycines* parasitism at a statistically significant level for the 15 MAPK-OE experiments overlap with the 14 MAPK-RNAi experiments. A comparison of statistically significant outcomes for the replicated OE and RNAi experiments have identified 9 MAPKs that are common between the MAPK-OE and MAPK-RNAi experiments (**Figure 3**). The combined observations of suppressed parasitism when overexpressing these MAPKs in the *H. glycines*-susceptible *G. max*_[Williams 82/PI 518671]_ and increased parasitism in RNAi experiments in the *H. glycines*-resistant *G. max*_[Peking/PI 548402]_ in both the wr and pg analyses strongly implicates the genes as having a defense role. To further place these results into context, analyses have been done to determine how many of these 9 MAPKs are expressed in the root cells prior to and/or at time points leading to a defense response. Analyses demonstrate that 12 MAPKs that are expressed within these syncytium cells undergoing their respective defense responses (**Supplemental Table 1**). Of these MAPKs, 7 have a defense role as has been indicated by the criteria set for the functional experiments (**Table 2**). These MAPKs include MAPK2, MAPK3-1, MAPK4-1, MAPK6-2, MAPK13-1, MAPK16-4 and MAPK20-2. Those MAPKs expressed within the syncytium, but not having a defense role include MAPK5-1, MAPK11-1, MAPK16-2, MAPK16-3 and MAPK19. None of the 9 MAPKs that do not measure expression within the syncytium have a defense role based on the set criteria (**Table 2; Supplemental Table 1**). Lastly, 2 of the 11 MAPKs whose expression could not be measured due to how the Affymetrix^®^ Gene Chip^®^ had been fabricated (MAPK3-2 and MAPK5-3) are shown here to have a defense role (**Table 2; Supplemental Table 1**). However, as noted earlier, MAPK3-2 and MAPK5-3 have been observed to be expressed in roots (Goodstein et al. 2012; Neupane et al. 2013). Experiments then have been designed to determine the gene expression that these different defense MAPKs have in relation to each other.

**Table 2.** Correlation between the expression of the MAPK and its role in defense. LCM-DCM either measured, did not measure or could not measure (no probe) MAPK gene expression within syncytia undergoing the process of defense in both the *G. max*_[Peking/PI 548402 + PI 88788]_ genotypes. A FI of OE and RNAi lines identified how those now proven defense MAPK genes relate to those three DCM bins (**Figure 3**).

| <u>LCM-DCM Outcome</u> | <u>Syncytium expressed</u> | <u>Defense role</u> | <u>Percent</u> |
| --- | --- | --- | --- |
| Measured | 12 | 7 | 58.3 |
| Not Measured | 9 | 0 | 0 |
| no probe | 11 | 2 | 18.2 |
| **Total** | 32 | 9 | n/a |

### *G. max* MAPKs function as part of a signaling cascade that leads to defense

Analyses having been performed in *G. max* show components of two different receptor systems, functioning in PTI and ETI, act in defense to *H. glycines* parasitism (Pant et al. 2014, 2015; McNeece et al. 2017; Aljaafri et al. 2017). Each of these receptor systems transduce signals through the MAPK cascade (Desikan et al. 2000; Zhang et al. 2011; Couto et al. 2016). Furthermore, EDS1 and LSD1 functions in *A. thaliana* at a hub, regulating immunity and suppressing the generation of reactive oxygen species produced in photorespiration (Mateo et al. 2004; Mühlenbock et al. 2008; Roberts et al. 2013). An examination of the gene expression of the 9 defense MAPKs is presented in relation to *G. max* treated with harpin (**Figure 4**). Furthermore, the influence of transgenic overexpressing and RNAi lines for NDR1-1, BIK1-6, EDS1-2, and LSD1-2 have been examined in relation to the 9 defense MAPKs (**Figure 4**).

**Figure 4.**
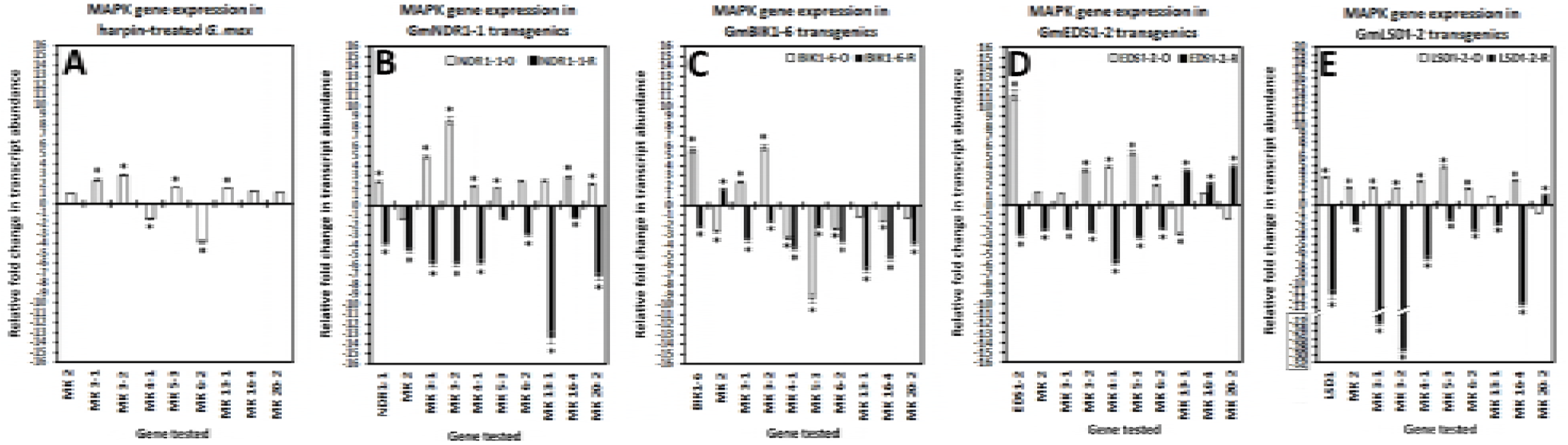
MAPK gene expression as measured by qPCR. **A**. harpin-treated *G. max*. **B**. NDR1-1 OE and RNAi transgenic lines. **C**. BIK1-6 OE and RNAi transgenic lines. **D**. EDS1-2-OE and RNAi transgenic lines. **E**. LSD1-2 OE and RNAi transgenic lines. A minimum cutoff of ± 1.5 is set. * statistically significant. The p-values for the replicated qPCR analyses have been calculated through a *t*-test (Yuan et al. 2006). Error bars represent standard deviation. Please refer to Methods for analysis details.

Harpin is a bacterial elicitor that activates a defense response with part of that response involving induced NDR1 gene expression (Wei et al. 1992; Gopalan et al. 1996). Harpin-treated *G. max* show increased relative transcript levels of MAPK3-1, MAPK3-2, MAPK5-3 and MAPK13-1 (**Figure 4**). In contrast, harpin treated *G. max* also show decreased relative transcript levels of MAPK4-1 and MAPK6-2 (**Figure 4**). No effect has been observed on the relative transcript abundance of MAPK2, MAPK16-4 and MAPK20-2 in harpin-treated *G. max* (**Figure 4**). These results show that harpin functions in specific ways in relation to MAPK gene expression.

NDR1 is a CC-NB-LRR protein known to function upstream of MAPK signaling and downstream of harpin treatment (Gopalan et al. 1996; Desikan et al. 1999). Furthermore, harpin treatment of *G. max* has been shown to increase the relative transcript abundance for NDR1-1 (Aljaafri et al. 2017). Analyses of *G. max* roots engineered to increase the relative transcript abundance of GmNDR1-1 by overexpression or decrease it by RNAi results in differential expression for a number of its MAPKs (**Figure 4**). Notably, MAPK3-1, MAPK3-2, MAPK16-4 and MAPK20-2 exhibit increased relative transcript abundance levels in the NDR1-1-OE lines. In contrast, MAPK3-1, MAPK3-2, MAPK16-4 and MAPK20-2 also exhibit decreased relative transcript abundance levels in the NDR1-1-RNAi lines (**Figure 4**).

BIK1 is a cytoplasmic receptor like kinase that has been proposed to function in *A. thaliana* through MAPK signaling (Veronese et al. 2006; Lin et al. 2014; Couto et al. 2016). Analyses of *G. max* roots engineered to increase the relative transcript abundance of BIK1-6 by overexpression or decrease it by RNAi results in differentially expressed gene expression of MAPK3-1 and MAPK3-2 (**Figure 4**). In contrast, MAPK2 exhibits decreased relative transcript abundance in the BIK1-6-OE lines while having increased relative transcript abundance in the BIK1-6-RNAi lines (**Figure 4**). The other MAPKs exhibit complex types of gene expression. These complex types of gene expression include decreased relative transcript abundance in both the BIK1-6 OE and RNAi lines (MAPK4-1, MAPK5-3, MAPK6-2 and MAPK16-4) and decreased relative transcript abundance in the RNAi lines with no effect observed in the OE lines.

EDS1 is a lipase that can receive input signals transduced from TIR-NB-LRRs such as RPP4 and DSC1 to engage defense responses (Aarts et al. 1998; Falk et al. 1999; van der Biezen 2002; Huang et al. 2010a). In the analysis presented here, *G. max* roots made transgenic to overexpress or undergo RNAi of EDS1-2 result in the differential expression of MAPK3-2, MAPK4-1, MAPK5-3 and MAPK6-2 (**Figure 4**). In contrast, MAPK13-1 exhibits a decrease in relative transcript abundance in the EDS1-2-OE lines (**Figure 4**). The remaining MAPKs exhibit no statistically significant change in expression in the EDS1-2-OE lines (**Figure 4**). While, MAPK2 and MAPK3-1 do not reflect an increase in relative transcript levels in the EDS1-2 OE lines, they do exhibit decreased transcript levels in the EDS1-2-RNAi lines (**Figure 4**).

LSD1 is a GATA-type β-ZIP transcription factor identified in *A. thaliana* functioning during a number of physiological processes (Dietrich et al. 1997; Wituszynska et al. 2013, 2015; Szechyńska-Hebda et al. 2016). The *G. max* LSD1-2 has been shown to affect the expression of many defense genes, including EDS1-2, involved in defense in the *G. max*-*H. glycines* pathosystem (Pant et al. 2015). Furthermore, overexpression of the *G. max* NDR1-1 results in induced LSD1-2 expression (McNeece et al. 2017). These observations are in agreement with the fundamental role that the *A. thaliana* LSD1 has in cell physiological processes (Mateo et al. 2004; Roberts et al. 2013). In the analysis presented here, *G. max* roots made transgenic to overexpress or undergo RNAi of LSD1-2 result in the differential expression of a number of defense MAPKs (**Figure 4**). In contrast, the relative transcript abundance of MAPK13-1 is not affected in the LSD1-2-OE lines while exhibiting decreased relative transcript abundance in the RNAi lines. MAPK20-2 exhibits no change in relative transcript abundance in the LSD1-2-OE lines while exhibiting an increase in relative transcript abundance in the RNAi lines. These results indicate that there are a number of MAPKs whose expression is under different types of transcriptional control by LSD1-2. Pant et al. (2015) has presented results showing the effect that LSD1-2 overexpression and its RNAi have on the relative transcript abundance of EDS1-2. However, results have not been presented showing the effect that EDS1-2 overexpression or RNAi have on the relative transcript abundance of LSD1-2. The effect that EDS1-2 overexpression or RNAi has on the relative transcript abundance of LSD1-2 are presented here (**Supplemental Figure 4**). An analysis has then been done to summarize the observed increased expression of MAPKs found in the harpin-treated plants as well as NDR1-1, BIK1-6, EDS1-2 and LSD1-2 OE and RNAi lines (**Supplemental Table 4**). In other experimental systems, harpin, NDR1, BIK1 and EDS1 and LSD1 have been determined to function very early in defense signaling to control downstream events (Century et al. 1997; Desikan et al. 1999; Mateo et al. 2004; Veronese et al. 2006; Wu et al. 2011). Experiments presented here have then been designed to determine how a panel of proven defense genes functioning in the *G. max*-*H. glycines* pathosystem respond to the overexpression or RNAi of these MAPK defense genes. A summary of these genes is provided (**Supplemental Table 5**).

### A MAPK can influence the expression of other MAPKs

The FI results show 9 MAPKs have met the criteria set for them having a role in defense. Furthermore, 7 of these defense MAPKs have been observed to be expressed during the defense process occurring within the syncytium in two different *H. glycines*-resistant *G. max* genotypes. In contrast, the remaining two MAPKs lack the fabrication of probe sets on the Affymetrix^®^ array that otherwise may have allowed for the detection of their gene expression during the defense response under study. However, experiments have shown that they are expressed within the root (Goodstein et al. 2012; Neupane et al. 2013). RNA from those 9 MAPK-OE and RNAi lines have been used in two different, but interrelated analyses. The first analysis has determined whether the MAPKs exert any influence on each other’s expression (**Figure 5**). The analysis has then determined whether the expression of the MAPKs affects the transcription of proven defense components including upstream signaling processes involving harpin and NDR1-1 and genes proven to function in defense in the *H. glycines*-*G. max* pathosystem (**Figure 5**).

**Figure 5.**
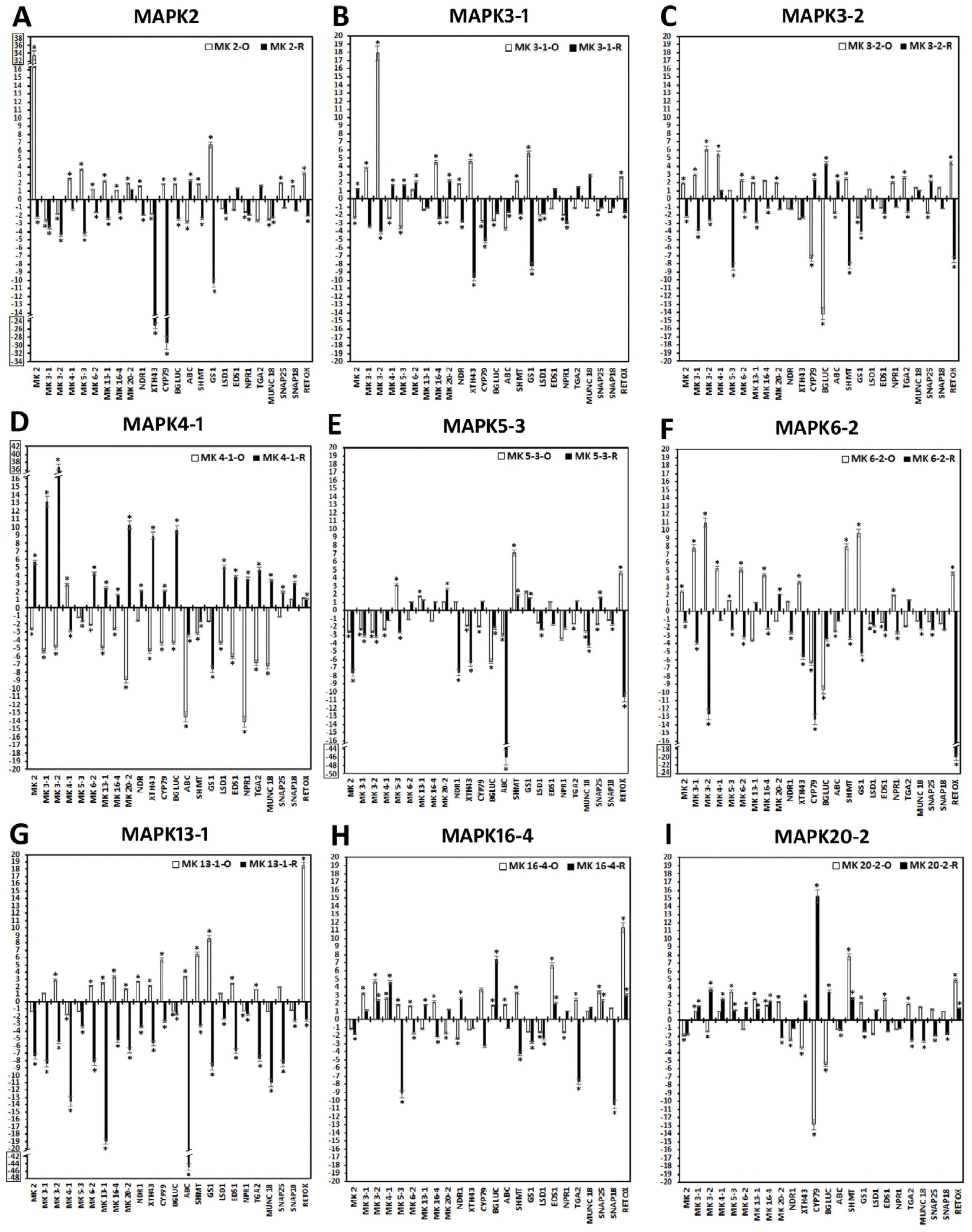
Coordinated gene expression among the defense MAPKs as measured by qPCR. **A**-**I**, gene expression among the MAPKs functioning in defense, including proven defense genes. A minimum cutoff of ± 1.5 is set. * statistically significant. Please refer to Methods for analysis details.

The experiments have identified five different types of differential expression that relate to the MAPKs (**Figure 6; Supplemental Figure 5; Supplemental Table 6**). Type 1 gene expression is characterized by the gene being induced in the OE treatment and also suppressed in the RNAi treatment or suppressed in the OE treatment and also induced in the RNAi treatment while in contrast, the gene is suppressed in the RNAi treatment and induced in the OE treatment or is induced in the RNAi treatment and also suppressed in the OE treatment (**Figure 6**). Type 2 gene expression is characterized by the gene being induced in the OE treatment and not differentially expressed in RNAi treatment or is suppressed in the OE treatment and not differentially expressed in RNAi treatment while in contrast, the gene is suppressed in the RNAi treatment and not differentially expressed in the OE treatment or is induced in the RNAi treatment and not differentially expressed in the OE treatment (**Figure 6**). Type 3 gene expression is characterized by the gene being induced or suppressed in co-regulated gene pair while in contrast, the gene is suppressed or induced in the co-regulated gene pair (**Figure 6**). Type 4 gene expression is characterized by the gene being induced and suppressed in the co-regulated gene pair and is also suppressed and induced in the co-regulated gene pair in the RNAi treatment while in contrast the gene is suppressed and induced in the co-regulated gene pair and is also induced and suppressed in the co-regulated gene pair in the OE treatment (**Figure 6**). The results demonstrate that while some overlap of gene expression occurs between some of the different MAPKs, each MAPK appears to have its own signature regarding the defense response and it is possibly this coordinated expression that leads to effective defense response. Type 5 gene expression is characterized by the gene being induced in both the OE and RNAi treatments within the gene pair or suppressed in OE and RNAi treatments within the gene pair (**Supplemental Figure 5; Supplemental Table 6**). A curious discovery in these gene expression studies has been that MAPK4-1 functions effectively in defense when overexpressed, but exhibits a decrease in many proven defense genes. This observation has merited additional study.

**Figure 6.**
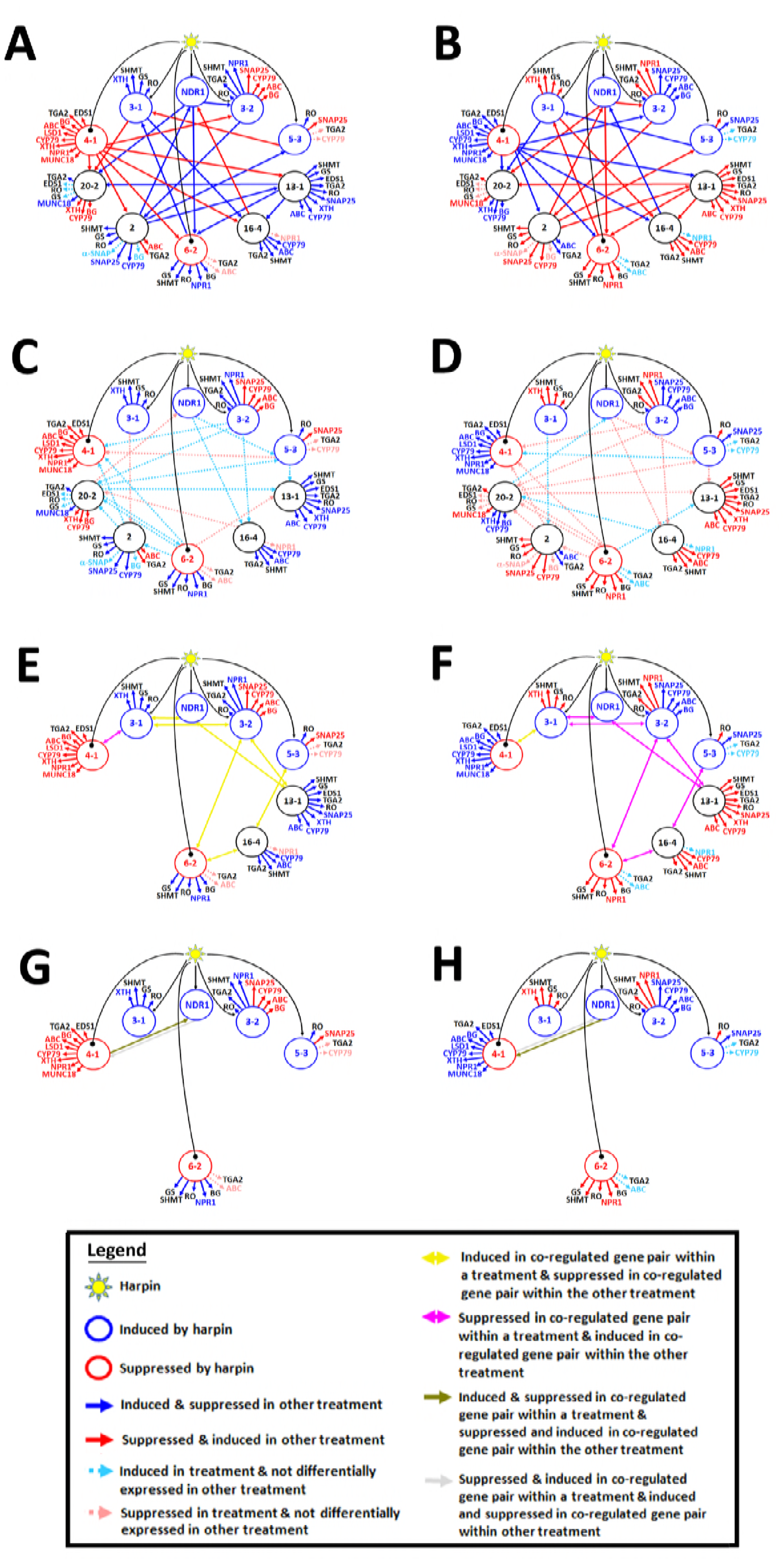
Graphic of coordinated gene expression presented in **Figure 5**. Type 1; **C**, **D**, Type 2; **E**, **F**, Type 3; **G**, **H**, Type 4; **I**, **J**. **A**, Type 1-OE: induced in the OE treatment and also suppressed in the RNAi treatment or suppressed in the OE treatment and also induced in the RNAi treatment. **B**, Type 1-RNAi: suppressed in the RNAi treatment and induced in the OE treatment or induced in the RNAi treatment and also suppressed in the OE treatment. **C**, Type 2-OE: induced in the OE treatment and not differentially expressed in RNAi treatment or suppressed in the OE treatment and not differentially expressed in RNAi treatment. **D**, Type 2-RNAi: suppressed in the RNAi treatment and not differentially expressed in the OE treatment or induced in the RNAi treatment and not differentially expressed in the OE treatment. **E**, Type 3-OE: induced or suppressed in co-regulated pair. **F**, Type 3-RNAi: suppressed or induced in co-regulated pair. **G**, Type 4-OE: Induced and suppressed in co-regulated pair and also suppressed and induced in co-regulated pair in the RNAi treatment. **H**, Type 4-RNAi: suppressed and induced in co-regulated pair and also induced and suppressed in co-regulated pair in the OE treatment. Please refer to legend for details (**Supplemental Table 6**). Please refer to Methods for analysis details. Harpin and Type 5 expression are presented (**Supplemental Figure 5**).

### Determining how GmMAPK4-1 could function in defense

While the observation that the overexpression of GmMAPK4-1 leads to suppressed gene activity for each of the proven defense genes under study while still functioning in defense is curious. However, MAPK4 genes have been shown to function in defense in different plant pathosystems (Zhou et al. 1999; Andreasson et al. 2005; Qiu et al. 2008; Zhang et al. 2016). These observations indicate the presence of different proteins, perhaps, functioning uniquely to various pathogens like *H. glycines*. Prior observations have identified increased expression of a *G. max PR1* homolog (GmPR1-6) (Glyma15g06780) by GmNPR1 (Pant et al. 2014). GmPR1-6 is part of a family of 9 related genes found at two loci on chromosomes 13 and 15 and its expression is shown here to occur in syncytia undergoing the process of defense (**Supplemental Table 7**). Experiments presented here show PR1-6 is induced in its expression in each of the defense MAPK lines while being suppressed in each RNAi line, including GmMAPK4-1 (**Supplemental Figure 6**). Furthermore, PR1-6 has characteristics of a secreted protein, including a predicted signal peptide (**Supplemental Figure 7**). These observations have led to functional experiments aiming to determine if PR1-6 has a defense function toward *H. glycines* parasitism. qPCR studies show PR1-6 expression is increased in each of the PR1-6-OE lines while being decreased in the PR1-6-RNAi lines (**Supplemental Figure 8**). Functional overexpression and RNAi transgenic analyses of PR1-6 shows it has a defense role (**Figure 7**). The results have identified that PR1-6 is a GmMAPK4-1 induced gene, along with being induced in each of the other defense MAPKs and functions in defense. Consequently, the results clarify how GmMAPK4-1 could function in the defense process.

**Figure 7.**
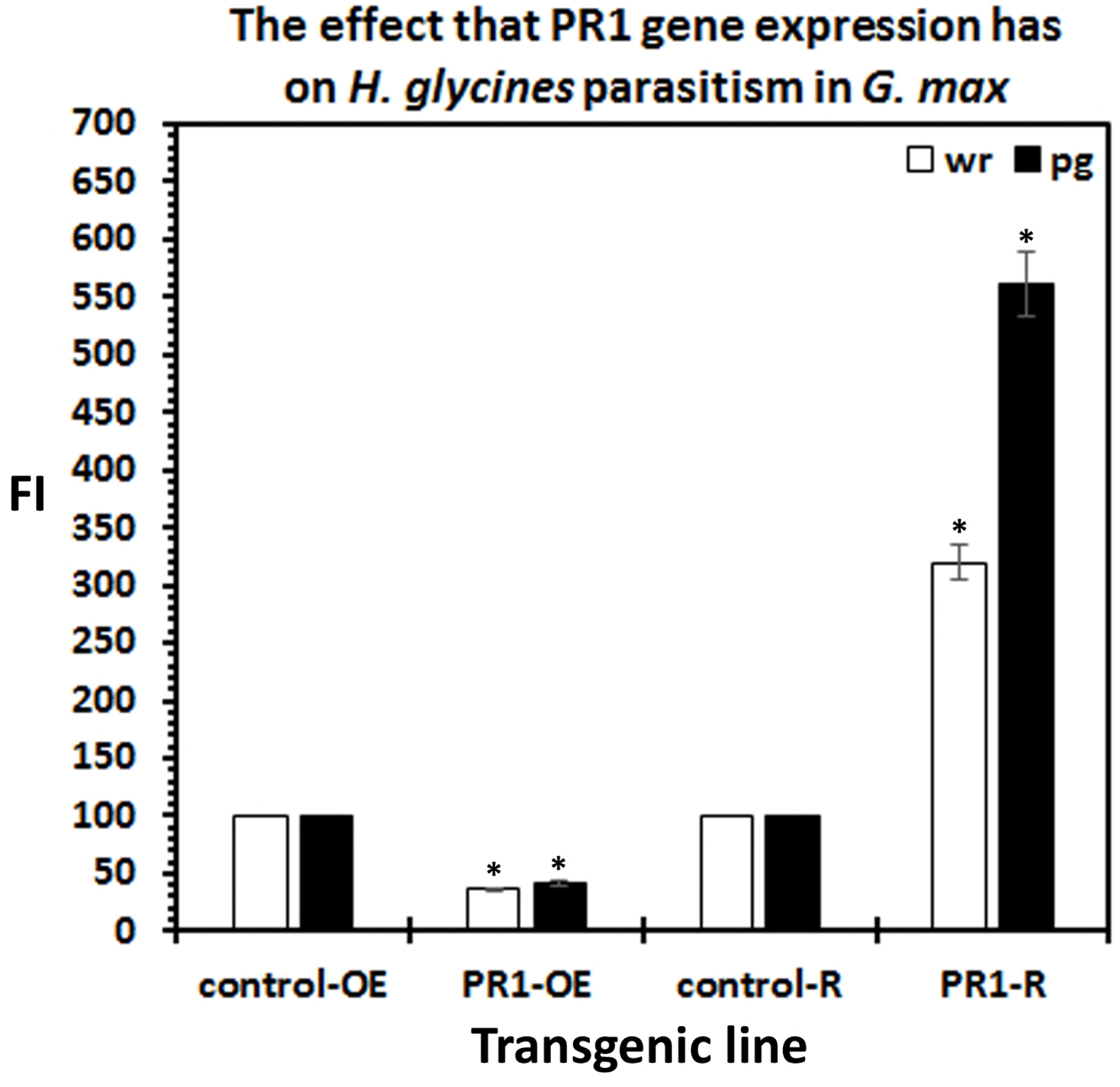
PR1-6 functions in defense to *H. glycines*. PR1-6 (Glyma15g06780, GmaAffx.36484.1.S1_s_at) is expressed specifically in syncytia undergoing the process of defense as revealed by DCM (**Supplemental Table 7**). **A**. The relative transcript abundance of the PR1 gene has been either increased through overexpression (OE) or decreased through RNAi (R). **B**. The calculated FI for the OE and RNAi lines as compared to the controls; p < 0.05 calculated by the Mann–Whitney–Wilcoxon (MWW) Rank-Sum Test which is a nonparametric test of the null hypothesis not requiring the assumption of normal distributions (Mann and Whitney, 1947). The experimental error representing standard deviation is presented. **C**. PR1-6 is induced in its expression in the MAPK OE lines and suppressed in its expression in the RNAi lines, p < 0.05.

## DISCUSSION

The study presented here using *G. max* as a model is a functional analysis of an entire MAPK gene family. Using this model, the analysis has identified MAPK gene expression occurring within the pathogen-parasitized root cells of two different *G. max* genotypes that undergo different, but related, forms of the defense process (Caldwell et al. 1960; Endo 1965, 1991; Klink et al. 2011; Matsye et al. 2011). The functional work has transgenically-altered the transcription of each MAPK family member allowing for the determination of its relationship to infection by a root pathogen and PTI and ETI signaling (**Figure 8**).

**Figure 8.**
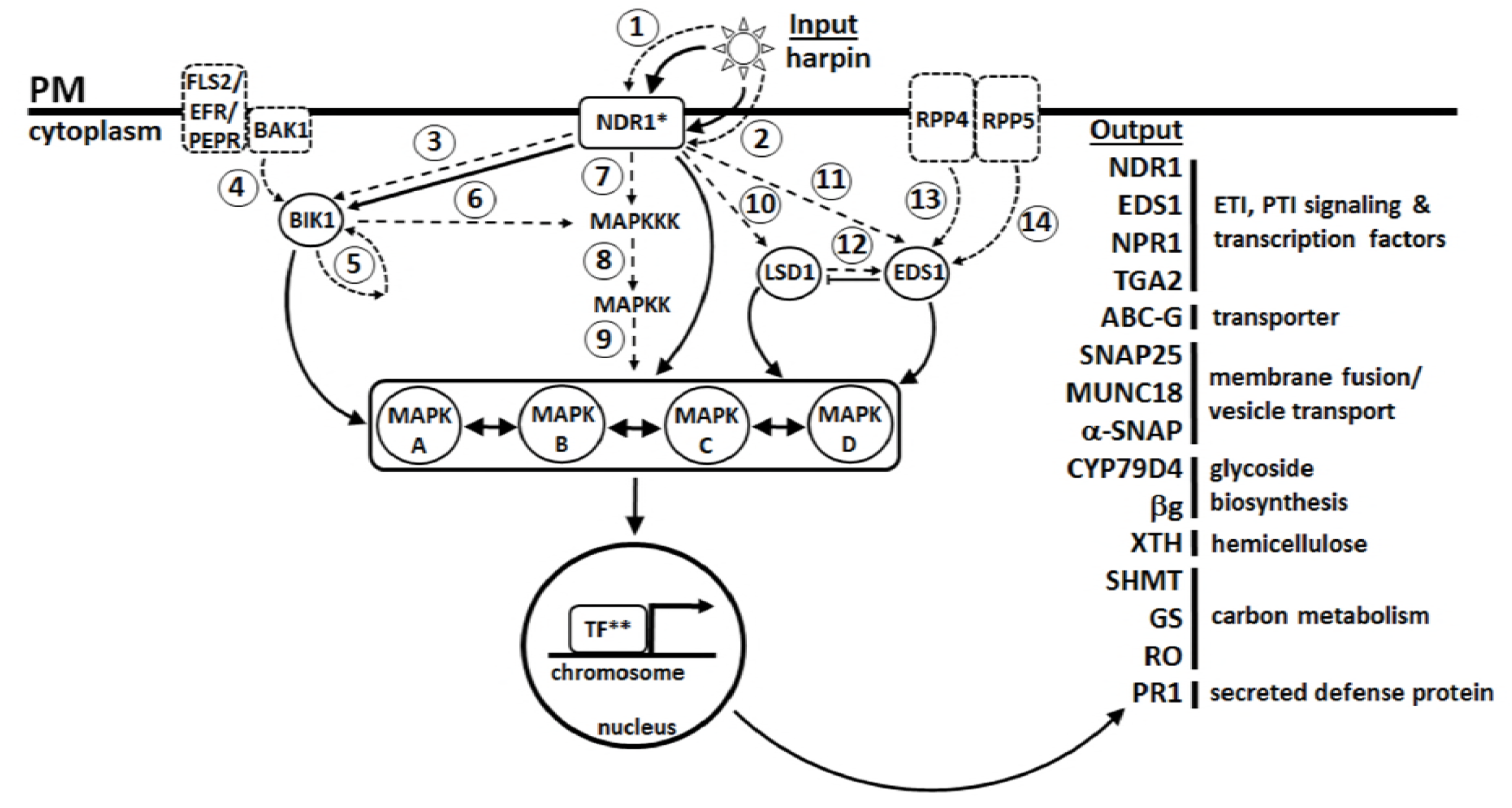
Model. The harpin effector acts as an input either outside of the plasma membrane through tissue infiltration or other means (**1**) or secreted inside of the cell through a type III secretion system (**2**). The result increases the relative transcript abundance of NDR1 (*). NDR1 activates BIK1 (**3**) which is released from BAK1 by phosphorylation (**4**) or can be autophosphorylated (**5**) likely activates MAPK signaling, a module identified to function in plant defense (**6**). Harpin probably activates MAPK signaling through NDR1 (**7**). Signal transduction proceeds through MAPKK (**8**) and MAPK (**9**) tiers. The experiments show that the expression of the four different types of MAPKs (A-D) is increased by NDR1-1 overexpression and decreased by its RNAi. Cross-talk occurs between the different MAPKS (↔). MAPKs influence the expression of defense TFs** such as NPR1-2, TGA2-1 and LSD1-2. The output of this MAPK module is shown has a number of different proven defense genes that are involved in a coordinated effort to stave off parasitism. References (**1**) Wei and Beer, 1993; Charkowski et al. 1998, (**2**) Wei and Beer, 1993, (**3**) Chang and Nick 2012, (**4**) Li et al. 2002; Chinchilla et al. 2007; Lu et al. 2010a, b, Wu et al. 2011; (**5**) Lin et al. 2014 (**6**) Zhou et al. 2014, Asai et al. 2002 (**7**) Desikan et al. 2001; Lee et al. 2001, (**8**), Anderson et al. 1990; Zhang and Klessig 2001; Jonak et al. 2002; (**9**) Sturgill and Ray, 1986; Zhang and Klessig 2001; Jonak et al. 2002; (**10**) Aljaafri et al. 2017; (**11**) Aljaafri et al. 2017; (**12**) Pant et al. 2015; (**13**) Aarts et al. 1998; (**14**) Aarts et al. 1998.*Gopalan et al. 1996. **Pant et al. 2014, 2015. Processes demonstrated by other cited authors are presented as dashed lines. Processes presented as solid lines are demonstrated in this analysis, McNeece et al. 2017 or Aljaafri et al. 2017. The output genes have been presented (Matsye et al. 2012; Pant et al. 2014, 2015; Sharma et al. 2016; McNeece et al. 2017; Klink et al. 2017). FLS2/EFR/PEPR and BAK1 are delimited by a dashed line since they have not been shown to be involved in *G. max* defense to *H. glycines*, but represent receptors that activate ETI and PTI, respectively. GmBIK1-6 and GmLSD1-2 expression are not induced as an output by the MAPKs under study under the analysis parameters.

### MAPKs expressed within the syncytium function during the defense process in the root

The experiments presented here show that MAPK gene expression found at the site of a resistant reaction is a good indicator that the gene functions in defense in the *G. max*-*H. glycines* pathosystem. Therefore, the results further confirm the hypothesis that gene expression occurring at the site of infection during defense function in the defense process (Klink et al. 2007; Matsye et al. 2011, 2012). Furthermore, the results indicate that the gene could function in resistance even if a defense role for the gene previously had not been known (Matsye et al. 2012; Matthews et al. 2013; Pant et al. 2014, 2015; Sharma et al. 2016; McNeece et al. 2017; Klink et al. 2017). Therefore, the platform presented here provides a general framework to perform those analyses.

Functional studies have identified a subset of 9 of the 32 *G. max* MAPKs that function effectively in defense, based on the study criteria. Within the cluster of defense MAPKs are *G. max* homologs of *A. thaliana* MPK2, MPK3, MPK4 and MPK6. Consequently, the procedures have led to the identification of the types of MAPKs that are expected to be functioning in defense processes, including those found in *A. thaliana* to function in defense to parasitic nematodes (Petersen et al. 2000; Desikan et al. 2000; Yuasa et al. 2001; Asai et al. 2001; Brodersen et al. 2006; Su et al. 2007; Qiu et al. 2008; Sidonskaya et al. 2016). In contrast, a number of defense MAPKs have been identified here that had not yet been shown through functional experiments to have a defense role. These MAPKs include *G. max* homologs of MPK5, MPK13, MPK16 and MPK20. Of these MAPKs, only *A. thaliana* MPK5 in has been shown to be associated with a process (salt stress) (Nühse et al. 2000). However, this result has not been tested functionally in *A. thaliana*. Furthermore, *A. thaliana* MPK13 has been shown to be activated by elicitor treatment, although a functional role has not been determined (Nitta et al. 2014). The functional experiments presented here, along with the accompanying gene expression work, has provided a characterization. Subsequent studies have aided in understanding their regulation further.

Some MAPKs that have been observed to be expressed within the syncytium undergoing the defense response did not meet the set criteria for functioning in defense. Altered MAPK signaling is known to activate autoimmune responses, leading to affected development (Petersen et al. 2000; Jammes et al. 2009; Kosetsu et al. 2010; Bethke et al. 2012; López-Bucio et al. 2014; Lang et al. 2017). Consequently, in the experiments presented here it is possible to have a reduced FI value while analyzing parasitism with regard to the number of cysts found on the whole root which commonly is done in this pathosystem. However, in these cases, one possible reason why the reduction in FI is observed could be because there is a much smaller root mass for the nematode to parasitize. Consequently, the FI would not be statistically significantly lower in this case in the pg analyses. For example, under the set criteria, MAPK5-1 is expressed in syncytia during a defense reaction at the 6 dpi time point. In the MAPK5-1-OE lines, a statistically significantly lower amount of *H. glycines* parasitism has been identified in the wr analysis, (FI = 37.6, SD = 9.6) that had not been observed in the pg analyses (FI = 67.7, SD 17.7). Furthermore, in the RNAi lines, no statistically significant increase in parasitism has been observed in the wr analyses (FI = 212.9, SD = 104.6), although a statistically significant increased level of parasitism has been observed in the pg analyses (FI = 221.4, SD = 31.0). Similarly, MAPK11-1 is expressed within the syncytium undergoing defense. MAPK11-1-OE lines did not exhibit statistically significant reduction in *H. glycines* parasitism in the wr or pg. Furthermore, MAPK11-1-RNAi lines do not meet the minimum threshold for an increase in parasitism in the wr analysis, but do in the pg analysis. In *A. thaliana*, flg22, bacterial elongation factor EF-Tu (elf18) and fungal N-acetylchitooctaose (ch8) elicitors can increase the expression and activate MPK11, indicating it functions in defense (Bethke et al. 2012; Eschen-Lippold et al. 2012). However, *mpk11* mutants fail to increase susceptibility to a number of pathogens including, *Alternaria brassicicola*, *Botrytis cinerea*, or *Pseudomonas syringae* (Bethke et al. 2012). In contrast, *mpk11* mutants are more susceptible to the plum pox potyvirus (Carrasco et al. 2014). The results demonstrate that the effects that the different MAPKs exert on pathogens may be very specific and the outcome may contrast based on the pathogen understudy. The results presented here show such an outcome occurs for the MAPK16 clade of parologous genes. While MAPK16-2, MAPK16-3 and MAPK16-4 are expressed within syncytia undergoing defense, only MAPK16-4 met the set defense criteria. For MAPK19-OE only the wr analysis met the set defense criteria, while the RNAi lines met the set criteria in the wr and pg analyses. No published functional work has been done in *A. thaliana* regarding MPK16 or MPK19 so the information generated here is new.

### Elicitor and receptor signaling lead to the induction of MAPK gene expression

The experiments presented here have revealed that the harpin treatment and the genetically engineered NDR1-1, BIK1-6, EDS1-2 and LSD1-2 lines all consistently affect the expression of the MAPK3-2 paralog. Other studies have shown that an increase in MAPK gene expression results in an increase in MAPK transcript, protein content and phosphorylation so the results presented here are expected to reflect their biological function (Xiong and Yang, 2003; Ren et al. 2006; Wang et al. 2016). Consequently, the results presented here indicate that both PTI and ETI branches of defense signaling converge on MAPK3-2, consistent with findings presented in other pathosystems (Wei et al. 1992; Century et al. 1995, 1997; Gopalan et al. 1996; Dittrich et al. 1997; Asai et al. 2002; Shoresh et al. 2006; Veronese et al. 2006; Kandoth et al. 2007; Wang et al. 2013; Lin et al. 2014; Couto et al. 2014). However, the experiments presented here have shown that only the MAPK3-2 paralog has its expression increased by NDR1-2 and EDS1-2. The results indicate the two *G. max* MAPK3 paralogs are under different types of regulation while some aspects appear to be coordinated and co-regulated. This topic will be discussed in a later paragraph.

The studied PTI and ETI components affect defense MAPK gene expression in different ways. For example, LSD1-2 has been shown here to induce the expression of the most defense MAPKs. The *A. thaliana* LSD1 has been shown to influence many physiological processes, consistent with the results presented here and elsewhere (Wituszynska et al. 2013, 2015; Pant et al. 2015). In contrast, altered BIK1-6 expression influences the fewest, affecting each of the two MAPK3 paralogs. Many studies presented in *A. thaliana* and other plant pathosystems have shown MPK3 (MAPK3) is central to many different types of defense responses (Asai et al. 2002; Gao et al. 2008). Harpin and NDR1-1 share the ability to influence the expression of MAPK3-1, MAPK3-2 and MAPK16-4, consistent with the demonstrated effect the elicitor has on NDR1 expression in different plants (Gopalan et al. 1996; Miao et al. 2010; McNeece et al. 2017). The results presented here indicate that harpin, NDR1-1, BIK1-6, EDS1-2 and LSD1-2 are capable of increasing the expression of different suites of MAPKs, possibly underlying the different types of resistant reactions that *G. max* has to *H. glycines* (Ross, 1958; Caldwell et al. 1960; Endo et al. 1965, 1991; Mahalingham and Skorupska 1996; Concibido et al. 2004; Ma et al. 2006; Klink et al. 2011; Pant et al. 2014, 2015). Alternatively, they function in concert to drive very effective defense processes found in various *G. max* genotypes.

### Harpin and NDR1 relate to MAPK signaling and defense responses in the root

Harpin has been reported to be responsible for the engagement of MAPK signaling processes through involving processes that increase the relative transcript levels of the NDR1 receptor (Gopalan et al. 1996; Zhang and Klessig 2000; Desikan et al. 1998, 1999, 2001; Samuel et al. 2005). However, it must be noted that NDR1 has not been shown to be that receptor of harpin. Harpin, instead has a broad impact on gene expression. For example, experiments that have been performed in *Gossypium hirsutum* (cotton) engineered to express the *Xanthomonas oryzae* pv. *Oryzae* harpin *hpa1Xoo* have led to the differential expression of 530 shoot genes (Miao et al. 2010). Among the 229 induced genes were the *G. hirsutum NDR1* (Gh-NDR1) and MPK7 (CM081B01) (Miao et al. 2010). Consequently, the involvement of harpin during *G. max* defense to different parasitic nematodes, including *H. glycines*, *Meloidogyne incognita* and *Rotylenchulus reniformis*, is consistent with its important role in defense signaling (Gopalan et al. 1996; Aljaafri et al. 2017).

In *G. max*, harpin induces NDR1-1 expression as well as a number of defense genes, including the major resistance gene *rhg1* (α-SNAP-5), *Rhg4* serine hydroxymethyltransferase (SHMT-5), EDS1-2, TGA2-1, galactinol synthase (GS-3), reticuline oxidase (RO-40) and xyloglucan endotransglycosylase (XTH-43) (Matsye et al. 2011, 2012; Pant et al. 2014; Sharma et al. 2016; Klink et al. 2017; Aljaafri et al. 2017). The results indicate the ETI-engaging harpin can serve a prominent defense role in this pathosystem. However, the relationship between harpin and MAPK signaling in *G. max* has not been examined until this study. Experiments presented here show that harpin treatment results in the induction of a subset of MAPKs, including MAPK3-1, MAPK3-2, MAPK5-3 and MAPK13-1. As already stated, *A. thaliana* MPK3 has been shown to be involved in multiple types of defense responses, functioning in different plant species as well (Gao et al. 2008). MPK3 also has functions that are redundant with MPK6, meaning MPK3 can function in the absence of MPK6 (Han et al. 2010). Constitutively active *A. thaliana* MPK3 drives defense responses through the CC-NB-LRR SUPPRESSOR OF MKK1 MKK2 1 (SUMM2) (Genot et al. 2017). Currently, very little functional work has been done in *A. thaliana* regarding how activated MPK5 or MPK13 function in *A. thaliana*.

In contrast to the aforementioned results, harpin treatment has been shown here to decrease the expression of MAPK4-1 and MAPK6-2. These observations are notable as it relates to a series of experiments that have been performed in *A. thaliana* demonstrating harpin treatment has no effect on MPK4 and MPK6 gene expression (Desikan et al. 1998, 1999, 2001). The effect that harpin has on the transcription of *G. max* MAPK4-2 and MAPK6-1 paralogs has not been ascertained since they do not function as defense MAPKs so it is possible that their expression is not influenced by harpin. A series of inhibitor studies have demonstrated that there are at least two defense branches that are activated by harpin through phosphorylation (Desikan et al. 2001). One branch leads to the production of an oxidative burst (Desikan et al. 2001). A second branch leads to the activation of MPK4 and MPK6 by phosphorylation (Desikan et al. 2001). Regarding gene expression, neither MPK4 nor MPK6 transcription is altered by harpin treatment and other *A. thaliana* defense MAPKs such as MPK3 can substitute in the *mpk3* genetic background (Desikan et al. 2001; Han et al. 2010). Consequently, phosphorylation of these MAPKs is responsible for signal transduction in the *A. thaliana* system (Desikan et al. 2001).

In the experiments presented here, the suppression of MAPK4-1 transcription by harpin treatment is intriguing since a number of observations made in *A. thaliana* have indicated that MPK4 is a negative regulator of SA signaling and systemic acquired resistance, occurring in the nucleus by its release from a target protein (Petersen et al. 2000; Qiu et al. 2008). In *A. thaliana*, after flg22 elicitor treatment, phosphorylated MPK4 regulates transcription of the cytochrome P450 monooxygenase PHYTOALEXIN DEFICIENT3 (PAD3) through MAP Kinase substrate 1 (MKS) and interactions with the WRKY33 transcription factor (Qiu et al. 2008). Consequently, the loss of MAPK4-1 would be expected to promote a defense response. Our observations have identified the decreased expression of a number of important defense genes in MAPK4-1 overexpressing lines while those lines still produce a defense response. The result indicates that it is possible that different sets of yet to be identified defense genes are expressed in the MAPK4-1 overexpressing lines that lead to a successful defense response. As demonstrated here, one of those genes could be GmPR1-6. It is possible that this defense response is fundamentally different than the one observed in *A. thaliana mpk4* mutants. For example, the overexpression of *Brassica napus* MPK4 results in resistance to *Sclerotinia sclerotiorum* and *Botrytis cinerea*. Consequently, overexpression of MAPK4 genes are capable of inducing a defense response. Similar results have been observed in *Nicotiana tabacum*. The overexpression of its MPK4 homolog NtMPK2 results in resistance to *Pseudomonas syringae* (Zhang et al. 2016). More detailed analyses have identified how MPK4 could be activating defenses. For example, MPK4 has been shown to function in defense through MAP kinase 4 substrate 1 (MKS) and WRKY33 to transcribe *PAD3* which leads to camalexin production (Zhou et al. 1999; Andreasson et al. 2005; Qiu et al. 2008). Consequently, *MPK4* and *mpk4* can each function in defense in different contexts. Notably, over 400 proteins become more abundant in *mpk4* plants (Takáč et al. 2016). Furthermore, inactivation of MPK4 by HopAI1 results in activation of SUMM2-mediated defense responses, a protective outcome (Zhang et al. 2012).

Harpin also has been shown here to suppress MAPK6-2 expression. *A. thaliana* MPK3 and MPK6 are parologous genes having overlapping functions. The infection of *A. thaliana mpk3* or *mpk6* mutants by *Botrytis cinerea*, results in no decrease in ethylene production which accompanies a susceptible reaction. In complimentary studies, *mpk3*/*mpk6* double mutants abolished *Botrytis cinerea*-induced ethylene production in *A. thaliana* (Han et al. 2010). This outcome happens because the normal functions of MPK3 and MPK6 are to stabilize ACC synthase (ACS) through phosphorylation (Han et al. 2010). Consequently, the MPK3 paralog has been shown to be sufficient to compliment *mpk6* (Han et al. 2010). Transcriptomic analyses of *mpk3*, *mpk4* and *mpk6* plants, revealed a highly interwoven network of genes common between or unique to each background indicating each gene is important to plant defense (Galetti et al. 2011; Frei dit Frey et al. 2014).

### BIK1 engages the expression of MAPK paralogs functioning in defense

The *A. thaliana* BIK1 is an important receptor-like cytoplasmic kinase (RLCK) that functions at a key branch point occurring between brassinosteroid ligand-facilitated growth signaling mediated by BRASSINOSTEROID INSENSITIVE 1 (BRI1) and defense signaling (Li and Chory et al. 1997). The defense signaling branch which is mediated by several different microbe associated molecular pattern (MAMP) receptors including FLAGELLIN SENSING 2 (FLS2) and EF-Tu RECEPTOR (EFR) as well as the *Arabidopsis* DAMP PEPTIDE 1 RECEPTOR (AtPEPR1) are shared between BIK1 and BRI1-associated kinase 1 (BAK1) (Li and Chory, 1997; Veronese et al. 2006; Zipfel et al. 2006; Liu et al. 2013). In this receptor system, ligand (brassinolide)-bound-BRI1 is capable of directly phosphorylating BIK1, releasing it from the complex in the absence of BAK1, promoting growth processes (Nam et al. 2002; Lin et al. 2013). In contrast, ligand-activated FLS2 results in an instantaneous association occurring between FLS2 and BAK1 (Gómez-Gómez and Boller, 2000; Chinchilla et al. 2007; Heese et al. 2007). In one scenario, BIK1 then becomes phosphorylated, a process dependent on the FLS2 and BAK1 association (Lu et al. 2010a, b, Zhang et al. 2010). Upon phosphorylation, BIK1 disassociates from BAK1, activating defense responses that eventually involve the MAPK cascade (Li et al. 2002; Chinchilla et al. 2007; Lu et al. 2010a, b, Zhang et al. 2010, 2011; Wu et al. 2011; Schwessinger et al. 2011; Chang and Nick 2012). However, more recent studies have shown that in addition to transphosphorylation by BAK1 during defense responses, BIK1 can be autophosphorylated, indicating that it functions differently than BAK1 (Lin et al. 2014). Details of any relationship between BIK1 and MAPK have not been examined until this presented work, although the ability of MAPK to phosphorylate BIK1 have been inconclusive. In the results presented here, it is clear that BIK1-6 overexpression can increase the expression of MAPK3-1 and MAPK3-2 with the opposite outcome happening in the BIK1-6 RNAi lines.

BAK1 also interacts with proteins other than BIK1 to effect defense responses (Li et al. 2001; Nam et al. 2002; Chinchilla et al. 2007; Heese et al. 2007; Postel et al. 2010; Roux et al. 2011). These results indicate the defense roles that BAK1 performs may be broad. Like BAK1, BIK1 also interacts with a number of different proteins (Lu et al. 2010a, Zhang et al. 2010; Liu et al. 2013; Lin et al. 2013, 2014). While some of these proteins overlap with BAK1, BIK1 is uniquely involved in suppressing the growth processes that are induced by brassinolide perception by BRI1 (Lin et al. 2013). These observations indicate BIK1 activates different sets of defense responses than BAK1 in *A. thaliana*. For BIK1, part of these roles includes its function as a dual specificity kinase capable of activating different defense outcomes (Lin et al. 2014). For example, the overexpression of BIK1 in *A. thaliana* results in phosphorylation of Flg22-induced receptor-like kinase 1 (FRK1) which is a PTI marker (Lu et al. 2010a; Lin et al. 2014). This result is consistent with observations made in the *G. max*-*H. glycines* pathosystem with concomitant induced gene expression of the PTI gene NONEXPRESSOR of PR1 (NPR1) and the secreted hemicellulose modifying protein gene xyloglucan endotransglycosylase/hydrolase (XTH43) at 0 dpi (Pant et al. 2014). GmBIK1-6 is a highly expressed gene expressed specifically in syncytia undergoing a defense response (Matsye et al. 2011; Pant et al. 2014). In the experiments presented here, BIK1-6 overexpression leads to increased transcript levels of MAPK3-1 and MAPK3-2 while its RNAi leads to their reduced transcript levels. These observations are in agreement with the important defense role that MAPK3 has in many plant species. In contrast, the overexpression of GmBIK1-6 has the opposite effect on MAPK2, indicating a very different type of regulation exists between BIK1-6 and MAPK2. Complex forms of regulation of different groups of MAPKs have been observed as a consequence of pathogen infection (Dóczi et al. 2007). BIK1-6 overexpression exerts little to negligible effect on MAPK6-2, MAPK13-1, MAPK16-4 and MAPK20-2, in contrast to the RNAi results. These observations are in agreement of observations made in *A. thaliana* showing the induction of MPK3. However, in the results presented here MAPK6-2 expression is not induced in either the BIK1-6 overexpression or RNAi lines. Consequently, it is possible that the involvement of *A. thaliana* MPK6 in defense processes driven through BAK1 occur through these other cytoplasmic kinases (Gao et al. 2009; Schwessinger et al. 2011).

### EDS1 functions antagonistically with LSD1 to modulate defense responses

EDS1 and LSD1 function in defense (Dietrich et al. 1997; Falk et al. 1999; Muhlenbock et al. 2008). The *G. max* homologs of EDS1 and LSD1 each have been shown to function in the defense process that *G. max* has toward *H. glycines* parasitism (Pant et al. 2014, 2015). In *A. thaliana*, the LSD1 β-ZIP, GATA-type transcription factor plays an important role in growth and defense processes (Dietrich et al. 1994, 1997; Jabs et al., 1996; Kliebenstein et al., 1999; Rustérucci et al. 2001; Epple et al. 2001; Aviv et al. 2001; Mateo et al. 2004; Kaminaka et al. 2006; Muhlenbock et al. 2008; Huang et al. 2010b; Czarnoca et al. 2017; Matuszkiewicz et al. 2018). Experiments have also demonstrated roles in various root processes, including biotic and pathogenic interactions (Muhlenbock et al. 2007; Pant et al. 2015; Guan et al. 2016; Matuszkiewicz et al. 2018). LSD1 resides at a key position between redox changes occurring in plastoquinone pools caused by excess excitation energy (EEE) during photosynthesis and the suppression of EDS1/PAD4 and photorespiration processes (Muhlenbock et al. 2008). A decrease in reactive oxygen species and ethylene–induced programmed cell death occurs through the role that LSD1 has in increasing catalase and superoxide dismutase gene expression/activity (Jabs et al., 1996; Kliebenstein et al., 1999; Mateo et al. 2004; Huang et al. 2010b). The link of LSD1 to photorespiration is an intriguing one since SHMT functions in this process and in *G. max*, its SHMT-5 encodes the *Rhg4 H. glycines* resistance gene (Muhlenbock et al. 2008; Liu et al;. 2012; Matthews et al. 2013). While intriguing in its role in electron transport, the shift between oxidation/reduction of plastoquinone also has important ramifications in shoot to root signaling and it is this state that is important in activating LSD1 (Muhlenbock et al. 2008; Petrillo et al. 2014). In prior experiments the *G. max* LSD1-2 gene expression has been shown to be induced by NDR1-1 overexpression while also being reduced by its RNAi (McNeece et al. 2017). The genetic manipulation of *G. max* LSD1-2 has also been shown to result in altered transcription of a number of genes shown to have proven defense functions in the *G. max*-*H. glycines* pathosystem, including EDS1-2 and NPR1-2 (Pant et al. 2015). The demonstration in *G. max* that NDR1-2 overexpression leads to increased LSD1-2 transcription prompted experiments designed to determine how LSD1-2, itself, relates to MAPK gene expression. The experiments presented here show that the genetically engineered OE/RNAi of LSD1-2 results in the differential expression of a number of MAPKs. Importantly, *G. max* LSD1-2-OE results in the induction of its homologs of *A. thaliana* MPK2, MPK3, MPK4, MPK6, each having proven roles in various defense processes (Desikan et al. 2000; Petersen et al. 2000; Nühse et al. 2000; Asai et al. 2002). *G. max* LSD1-2-OE also results in the induced expression of its homologs of *A. thaliana* MPK5 and MPK16 that have no demonstrated defense function. Consequently, it appears that LSD1 may have broader roles in defense than known previously (Pant et al. 2015; Matuszkiewicz et al. 2018). Recent experiments performed in *A. thaliana* have indicated a mutation in *lsd1* results in impaired nematode parasitism (Matuszkiewicz et al. 2018). The experiments differ with a previously published role shown through OE and RNAi experiments that *G. max* LSD1-2 has in defense to *H. glycines* (Pant et al. 2015). The Pant et al. (2015) experiments have identified the expression of *G. max* LSD1-2 within nematode-induced feeding sites undergoing two different forms of the defense response caused when *G. max*_[Peking/PI 548402]_ and *G. max*_[PI 88788]_ are infected with *H. glycines*_[NL1-Rhg/HG-type 7/race 3]_. These results had been compared to outcomes performed in the same *G. max* genotypes, but infected with *H. glycines*_[TN8]_ which results in a susceptible reaction in each genotype. These types of cross comparative studies are not possible in the *A. thaliana*-*H. schachtii* pathosystem, made more complicated to perform due to the lack of R genes functioning against the nematode (Matuszkiewicz et al. 2018). Furthermore, the root mass of *A. thaliana* is comparably small which may complicate analyses of the absolute number of nematodes per gram of root tissue (Matuszkiewicz et al. 2018). Pant et al. (2015) analyzed the role of LSD1-2 through its overexpression in the susceptible genotype *G. max*_[Williams 82/PI 518671]_, resulting in nearly a 60% reduction in cyst production as revealed by the FI. There was little indication of deleterious effects on root growth due to LSD1-2-OE or LSD1-2-RNAi, since they appeared similar to controls and the percent difference in roots mass as compared to the control had been shown to be negligible (Pant et al. 2015). Consequently, the FI would be very similar to the published results (Pant et al. 2015). Since *G. max* has 4 LSD1-like genes, it is possible that the other copies of the gene may exhibit overlapping roles (Epple et al. 2003). MAPK signaling has not before been reported to be influenced in *lsd1* mutants (Wituszynska et al. 2013; Matuszkiewicz et al. 2018). In contrast to studies examining *lsd1* mutants, few studies have examined what happens when it or its paralogs are overexpressed. The overexpression of a rice LSD1 related gene results in plants that are more saline-alkaline (SA) stress resistant under NaHCO_3_ treatment, resulting in plants that had larger vegetative characteristic and accompanied by higher relative transcript levels of a number of genes that function to quell oxidative stress (Wang et al. 2005; Guan et al. 2016). Consequently, it appears that plant mass increases with the overexpression of LSD1.

### Parologous gene family members experience co-regulated expression

The *G. max* genome has undergone two duplication events, processes that impact MAPK gene number (Schmutz et al. 2010; Mohanta et al. 2015). A number MAPKs have duplicated copies within the *G. max* genome (Schmutz et al. 2010; Neupane et al. 2013; Mohanta et al. 2015. However, prior to these studies it has been unclear whether any influence at a functional level exists between parologous MAPKs gene family members on a pathogenic process or on the expression of these genes. We hypothesized that parologous MAPKs may indeed have influential properties regarding the transcription of parologous MAPKs or other MAPKs in general. This hypothesis has been based off of our prior work that has demonstrated how genes composing a complex defense structure called a regulon are co-regulated (Pant et al. 2014; Sharma et al. 2016; Klink et al. 2017). In the analysis presented here the hypothesis has been tested through the identification that *G. max* MAPK3-1 and MAPK3-2 are co-regulated. What is notable in our observations is that the expression of MAPK3-2 is induced by harpin, NDR1-1, BIK1-6, EDS1-2, LSD1-2 while MAPK3-1 is not induced by NDR1-1. The MAPK3-1 parolog, however, is induced by MAPK3-2 expression and vice versa. Regarding the study of large gene families in plants, the preponderance of work has been devoted to examining the expression of genes from large transcriptomic studies and determining whether the genes are co-expressed (Curtin et al. 2012; Tang et al. 2016; Zhang et al. 2016). These studies provide important insights, especially when attempting to evaluate the evolutionary history of the genes in duplicated genomes (Curtin et al. 2012). However, a functional analysis supported by altering the expression of the genes is relatively atypical. Such approaches have been shown to provide important understanding in biological processes in general, including those outside of the plant kingdom (Ohno 1970; Prince and Pickett 2002; Tate et al. 2006; Lynch and Conery 2006; Hittinger and Carroll 2007).

### MAPKs induce the gene expression of PR1, a secreted protein

Experiments have shown that NPR1 expression leads to the induction of PR1 (PR1-6) transcription. Experiments have shown that the overexpression of a number of genes functioning in defense in the *G. max*-*H. glycines* pathosystem induce PR1-6 gene expression (Pant et al. 2014). These genes include NPR1, α-SNAP-5, SYP38 and XTH (Pant et al. 2014). PR1-6 is a member of a family containing 9 PR-related genes. Furthermore, the PR1 genes are located on chromosomes 13 and 15 in regions that appear to have undergone localized duplications. Genes composing chromosomal loci containing duplicated gene family members have been shown to perform important roles during defense to parasitic nematodes (Cooke et al. 2012). These genes, including α-SNAP are known to perform important roles in secretion (Novick et al. 1980; Clary et al. 1990). During the course of the experiments presented here, it has been observed that experimentally induced transcription of GmMAPK4-1 leads to a successful defense response even though the expression of all of the examined proven defense genes had decreased. The result indicates that yet to be identified proteins may have defense roles. The experiments presented here show that PR1-6 is induced in its expression specifically within the syncytium undergoing a defense response. Furthermore, qPCR has demonstrated increased transcript levels of PR1-6 in each of the defense MAPK overexpressing lines, including GmMAPK4-1 with suppressed levels observed in the RNAi lines. Functional experiments have revealed PR1-6 functions in defense, consistent with observations described earlier showing MPK4 functions in defense (Zhou et al. 1999; Andreasson et al. 2005; Qiu et al. 2008; Berriri et al. 2012; Zhang et al. 2016). Furthermore, the observation strengthens previous work showing PR1 functions directly during defense responses (Shin et al. 2014; Chen et al. 2014; Chung et al. 2018; Nawrocka et al. 2018; Yang et al. 2018).

The analyses presented here has examined the entire MAPK gene family for an allotetraploid with regard to a defense process. The analysis has identified MAPKs that function in defense with some of them previously not shown to have defense functions. Some MAPKs that have not been shown in the *G. max*-*H. glycines* pathosystem to be expressed during the defense response appear to be able to contribute to defense process when their transcription is influenced ectopically. These observations indicate that synthetic forms of defense signaling may exist (Khakimov et al. 2015; Zheng et al. 2015). Consequently, their expression could be manipulated in ways that generate novel forms of defense that have not been observed or studies until now. These novel forms of defense identified through the experimental acquisition of gene function may allow for the identification of previously unknown cell signaling processes (Ohno 1970; Prince and Pickett 2002; Lynch and Conery 2006). Furthermore, the work may help provide insight or approaches into studying the seemingly competing processes of defense and symbiosis (Ryu et al. 2017).

## ACKNOWLEDGEMENTS

The authors are thankful for start-up support and teaching assistantships provided by the Department of Biological Sciences. The authors are thankful for an awarded competitive Special Research Initiative grant from the College of Arts and Sciences at Mississippi State University (VPK).

## SUPPLEMENTAL TABLES

**Supplemental Table 1.** LCM-DCM gene expression summary of studied MAPKs. The LCM-DCM analysis examined control (0 dpi pericycle), 3 and 6 dpi syncytia of *G. max*_[Peking/PI 548402]_ and *G. max*_[PI 88788]_ genotypes (*G. max*_[Peking/PI 548402 + PI 88788]_) undergoing a successful defense response (Klink et al. 2010). The plants used in the experiments to obtain and collect the sample sets for the Affymetrix^®^ hybridizations have been generated independently three different times (Klink et al. 2010). M, gene expression measured as determined by the set criteria having a p < 0.05 in all 6 arrays; N/M, gene expression not measured as determined by the set criteria since at least one of the 6 arrays have a p ≥ 0.05; n/a, not applicable since the Affymetrix^®^ microarray did not have a probe set fabricated for that gene. Please refer to Methods for analysis details.

**Supplemental Table 2.** The *G. max* Class E MAPKs not analyzed in the presented study because they are not related to MAPKs either structurally, functionally or phylogenetically.

**Supplemental Table 3.** PCR and qPCR primers used in the study.

**Supplemental Table 4**. A qPCR summary for harpin-treated *G. max* and transgenic NDR1-1, BIK1-6, EDS1-2 and LSD1-2. Y (yes), has a statistically significant effect on MAPK expression whereby OE and RNAi have the opposite effect on MAPK expression; N (no), no statistically significant effect on MAPK expression whereby OE and RNAi do not have the opposite effect on MAPK expression or no not meet the expression criteria of ± 1.5 fold, p < 0.05.

**Supplemental Table 5**. Defense gene summary. *Reference 1 is the primary reference used for the particular gene or treatment. **Reference 2 is the primary reference used for the demonstration the gene functions in defense in the *G. max*-*H. glycines* pathosystem.

**Supplemental Table 6.** Supplement to **Figure 6**: legend detail.

**Supplemental Table 7.** PR1 LCM-DCM analysis. Analysis methods have been performed as described previously. * The e-values have been obtained by blasting Glyma15g06780 (GmPR1-6) to the *G. max* proteome. The ordering of PR1 genes is based on chromosomal position chromosome1 →chromosome 20, explaining the order of the e-values.

## SUPPLEMENTAL FIGURES

**Supplemental Figure 1**. Studied *G. max* and *H. glycines* tissue. **A**. Transgenic root pRAP15 overexpression control prior to *H. glycines* infection, Bar = 1 cm; **B**. Transgenic root pRAP17 RNAi control prior to *H. glycines* infection, Bar = 1 cm. **C**. *H. glycines* cyst. Bar = 500 μm.

**Supplemental Figure 2**. MAPK matrix. **A**, MAPK overexpression **B**, MAPK RNAi; qPCR for each transgenic MAPK line is presented in order of the legend (**Supplemental Table 4**). Asterisk depicts the particular OE or RNAi line used to examine the relative transcript abundance of the other MAPKs by qPCR. Error bars represent standard deviation. Please refer to Methods for analysis details.

**Supplemental Figure 3**. The effect that transgenically-altered MAPK expression has on *G. max* root mass for MAPK overexpression and RNAi. Error bars represent standard deviation. * p < 0.05 calculated by the MWW Rank-Sum Test. Please refer to Methods for analysis details.

**Supplemental Figure 4.** The effect that EDS1-2 overexpression or EDS1-2 RNAi has on the relative transcript abundance of LSD1-2 relative transcript abundance. Error bars represent standard deviation. * p < 0.05 calculated by the MWW Rank-Sum Test. Please refer to Methods for analysis details.

**Supplemental Figure 5.** Gene expression Supplement to **Figure 6**. **A**. Harpin influences the expression of MAPKs. **B**. Type 5 gene expression.

**Supplemental Figure 6**. PR1 is increased in its expression in the MAPK OE lines and decreased in its expression in the RNAi lines, p < 0.05. The experimental error representing standard deviation is presented. Please refer to Methods for analysis details.

**Supplemental Figure 7**. PR1-6 signal peptide prediction using three different prediction programs. **A**. SignalP-4.1; **B**. Phobius; **C**. **P**rediSI.

**Supplemental Figure 8**. PR1-6 expression is increased the PR1-6-OE lines and decreased in its PR1-6-RNAi lines, p < 0.05. The experimental error representing standard deviation is presented. Please refer to Methods for analysis details.

## Abbreviations

MAPK: Mitogen activate protein kinase
(ETI): effector triggered immunity
(PAMP): pathogen activated molecular pattern
(PTI): triggered immunity

